# Quantifying habitat and landscape effects on composition and structure of plant-pollinator networks in the US Northern Great Plains

**DOI:** 10.1101/2021.02.12.431025

**Authors:** Isabela B. Vilella-Arnizaut, Henning Nottebrock, Charles B. Fenster

**Author notes:** Plant Ecology, Bayreuth Center of Ecology and Environmental Research (BayCEER), University of Bayreuth, Universitätsstr. 30, Bayreuth, Germany (Henning Nottebrock).

## Abstract

Community structure contributes to ecosystem persistence and stability. To understand the mechanisms underlying pollination and community stability of natural areas in a human influenced landscape, a better understanding of the interaction patterns between plants and pollinators in disturbed landscapes is needed. The Northern Great Plains still retain extensive tracts of remnant temperate grassland habitat within a matrix of varying land-uses. We used a network-based approach to quantify how temperate grassland attributes and landscape heterogeneity influence plant-pollinator community structure in natural habitats. We also quantified pollinator diversity and floral diversity to assess the functional role of temperate grassland attributes and the surrounding landscape on the composition of the plant-pollinator communities in natural habitats. We found that the amount of local nectar sugar and increased proportions of certain land-uses contribute to pollinator diversity that in turn influences the structure of interactions between plants and pollinators. Understanding the factors contributing to plant-pollinator network structure can guide management decisions to support resilient plant-pollinator communities and conserve the stability of pollination services.

## Introduction

Habitat loss and fragmentation are among the most prominent anthropogenic pressures impacting biodiversity globally (Foley et al. 2005; Tscharntke et al. 2005; Lundgren and Fausti 2015; Greer et al. 2016). Increased intensive agriculture has facilitated the conversion of natural vegetation to crop monoculture (Ramankutty and Foley 1999; Rashford et al. 2011; Wright and Wimberly 2013). Globally, temperate grasslands are considered to be at the greatest risk for biodiversity loss and ecosystem dysfunction, specifically due to extensive landscape conversion and low rates of habitat protection (Hoekstra et al. 2005). The disparity between landscape conversion and protection for grasslands is particularly concerning since grasslands represent essential habitat and resources for multiple species including insect pollinators. Animal-driven pollination is essential for the production of approximately 35% of crops worldwide (Klein et al. 2007; Vanbergen et al. 2013) and contributes to the reproduction of over 70% of flowering plant species (Potts et al. 2010). Nevertheless, there is an alarming global decline of insect pollinators largely attributed to anthropogenic pressures such as land-use intensification and widespread use of pesticides (Kearns et al. 1998; Steffan-Dewenter et al. 2005). Given these documented declines, there is an increased interest in conserving pollinator communities and their habitats to stabilize these critical ecosystem services and in turn, ecosystem function (Steffan-Dewenter et al. 2002; Kremen et al. 2002; Olesen et al. 2007; Peterson et al. 2010; Redhead et al. 2018; Jauker et al. 2019).

An understudied determinant of pollinator communities in natural areas is the diversity and configuration of the landscape that exists in the mosaic of a mixed-use agricultural and natural landscape. Pollinator communities are negatively impacted when their populations become increasingly isolated from valuable resources (i.e., nectar resources, pollen resources, nesting resources, etc.) (Olesen et al. 1994; Kearns et al. 1998; Garibaldi et al. 2011). Landscape composition (e.g., patch size) and configuration (e.g., fragmentation) can influence insect pollinator behavior and movement (Fahrig et al. 2011; Hadley and Betts 2012; Moreira et al. 2015; Sakai et al. 2016). For example, proximity to semi-natural habitats increases the abundance of both wild bees and honey bees in crops (Steffan-Dewenter et al. 2002; Kremen et al. 2004; Heard et al. 2007). Likewise, increased availability and diversity of flowering plants in natural areas benefits pollinator populations (Potts et al. 2003; Potts et al. 2005; Ponisio et al. 2019; Requier et al. 2020). Recent studies (Nottebrock et al. 2017) demonstrate that the level of available sugar resources to pollinators can have considerable effects on the outcome of plant-pollinator interactions. Thus, the consequences of landscape conversion could impact bee populations on numerous levels (i.e., hive survival and productivity) by decreasing availability and diversity of nutritional floral resources (Naug 2009; Pettis et al. 2013; Otto et al. 2016). Within the midwestern United States, the Northern Great Plains has served as a refuge for approximately 40% of the commercial honey bee colonies from May through October (USDA, 2014). The regional blooms provided by livestock-grazed pastures and grasslands in the Northern Great Plains have sustained transported honey bee colonies due to the presence and abundance of floral resources (Otto et al. 2016). However, honey bee colonies in the US and Europe continue to sustain annual losses which can be attributed to a combination of factors such as disease, pests, and pesticides (Williams 2002; Cox-Foster et al. 2007; Cox-Foster et al. 2009; Alaux et al. 2010; Spleen et al. 2013). These detrimental health factors may be due in part to landscape simplification limiting the abundance and diversity of floral resources available to pollinators (Tscharntke et al. 2005; Smart et al. 2016).

A second gap in the literature is the extent to which landscape use determines the ways in which pollinators and plants interact with one another. It is important to understand how interactions between plants and pollinators may be affected across spatial-temporal scales in order to preserve the stability of pollination services (Burkle and Alarcón 2011). While it is understood that land-use intensification negatively affects both plant and pollinator diversity (Vinson et al. 1993; Williams et al. 2002; Kremen et al. 2002; Potts et al. 2003; Knight et al. 2009; Potts et al. 2010; Garibaldi et al. 2011; Spiesman and Inouye 2013; Habel et al. 2019), relatively few studies have utilized a network-based approach to consider the ecological impacts of both landscape composition and configuration on plant-pollinator community structure and stability (Weiner et al. 2014; Moreira et al. 2015; Tylianakis and Morris 2017; Redhead et al. 2018; Jauker et al. 2019; Lazaro et al. 2020). In an ecological context, network theory has been utilized to examine how the mutualistic interactions within plant-pollinator communities influences their structure and in a broader sense, interpret the mechanisms behind biodiversity and community resilience (Memmott et al. 2004; Bascompte et al. 2006; Blüthgen et al. 2008; Dupont et al. 2009a; Hadley and Betts 2012; Spiesman and Inouye 2013; Soares et al. 2017; Redhead et al. 2018). Mutualistic networks, such as plant-pollinator networks, tend to exhibit structural patterns such as nestedness, which demonstrates a degree of interaction redundancy in the community and is associated with overall community stability (Jordano et al. 2003; Bascompte et al. 2003;2006). Using a network-based approach can help discern the overall structure of plant-pollinator communities embedded within varying disturbed landscapes and their response to resource availability across space and season.

We focus our study in the Northern Great Plains within a region in eastern South Dakota known as the Prairie Coteau. Historically, the Great Plains stretched across approximately 60 million hectares in North America, though presently approximately 4% of this temperate grassland biome remains undisturbed from cropland conversion (Bauman et al. 2016). South Dakota is a part of a greater region known as the Western Corn Belt where the rate of grassland conversion to corn and soy agriculture fields reached ~1.0-5.4% annually from 2006 to 2011 (Wright and Wimberly 2013). Even though agriculture remains a primary land-use in much of the Upper Midwest, the Prairie Coteau region is unique in that approximately 17% of its original temperate grassland cover remains intact (Bauman et al. 2016). This provides an ideal study region to investigate plant-pollinator communities in remnant habitats established within an actively transforming working landscape. Considering the alarming decline of insect pollinators and grassland communities, there is a need to understand the community-level impact of habitat attributes and the surrounding landscape within an increasingly disturbed environment.

Agriculture-dominated landscapes such as the Western Corn Belt will need to coexist with the remaining patches of conserved natural habitat in order to continue utilizing the ecosystem services these habitats provide. For example, oil and pulse crops (e.g., sunflower, canola, legumes), which consist of approximately 1.1% of the landcover in South Dakota in 2019 (Han et al. 2012), benefit from insect pollination through increased crop yield (Tamburini et al. 2017; Mallinger et al. 2019). Within the Northern Great Plains, insect pollination increases sunflower crops by up to 45%, translating to a regional economic value of $40 million (Mallinger et al. 2019). Even primarily autogamous soybean crops, benefit from insect pollination through increased crop yield (Chiari et al. 2005; Lautenbach et al. 2012; Milfont et al. 2013; Cunningham-Minnick et al. 2019). Soybean crops occupy 7.1% of the landcover in South Dakota and are the second largest agricultural crop in the state behind corn (landcover of 9.01%) (Han et al. 2012). The need to conserve stable plant-pollinator interactions extends beyond the boundaries of remnant natural habitats and understanding how compounding local and landscape variables influence their outcome should be prioritized.

Our overall goal is to understand how plant-pollinator community structure is influenced by factors acting at various spatial-temporal scales. Rather than focus on the contribution of the surrounding natural landscape on pollinator services in agricultural fields, we take a novel approach to address how a mixed-use landscape affects pollinators found in natural habitats. Our study is sequential. We quantify the effects of habitat attributes and the surrounding landscape on pollinator diversity and then examine the relationship between pollinator diversity and the overall structure of the plant-pollinator communities by addressing the following questions: 1) How do North American temperate grassland attributes (i.e., patch size, floral diversity, latitude, nectar resources) affect pollinator diversity? 2) Is there a relationship between landscape and pollinator diversity, and if so, at what scale? Furthermore, does the proportion of land-uses surrounding North American temperate grasslands influence pollinator diversity, and if so, at what scale? and 3) How does pollinator diversity influence the overall structure of plant-pollinator communities? Essentially, given the plant community present at a site, how does pollinator diversity affect the community structure of interactions? These questions become especially relevant considering landscape conversion is expected to continue to reduce the natural landscape (Benton et al. 2003; Bauman et al. 2016; Liu et al. 2019). A better understanding of plant-pollinator networks and how their structure may be impacted by habitat or landscape attributes can help future management decisions focus on strategies promoting resilient plant-pollinator communities and conserve the stability of pollination services.

## Materials and Methods

### Study area

Formed by glacial uplift, the rolling hills of the Prairie Coteau span roughly 810,000 hectares across southeast North Dakota, eastern South Dakota, and southwest Minnesota (Bauman et al. 2016). Within South Dakota, this region covers approximately 17 counties and contains a matrix of land uses ranging from undisturbed grasslands to intensive cropland. Approximately 66% of eastern South Dakota has some type of crop disturbance history (Bauman et al. 2016). Currently, undisturbed grassland under protection from future conversion represents about 4% of the total land base in eastern South Dakota (Bauman et al. 2016). Thus, the grasslands that remain are fragments nestled within an increasingly disturbed landscape. The Prairie Coteau region represents a valuable resource for remnant temperate grassland habitat considering 17% of undisturbed grassland in eastern South Dakota still remains intact (Bauman et al. 2016). We selected fifteen remnant temperate grassland sites within the Prairie Coteau based on size, local landscape use, and proximity to other semi-disturbed grasslands (Table 1). Sites ranged in size from 8 to > 400 hectares and are managed by the US Fish & Wildlife Service (2), The Nature Conservancy (5), South Dakota Game, Fish & Parks (5) and the city of Brookings (1) with varied management regimes.

**Table 1.**
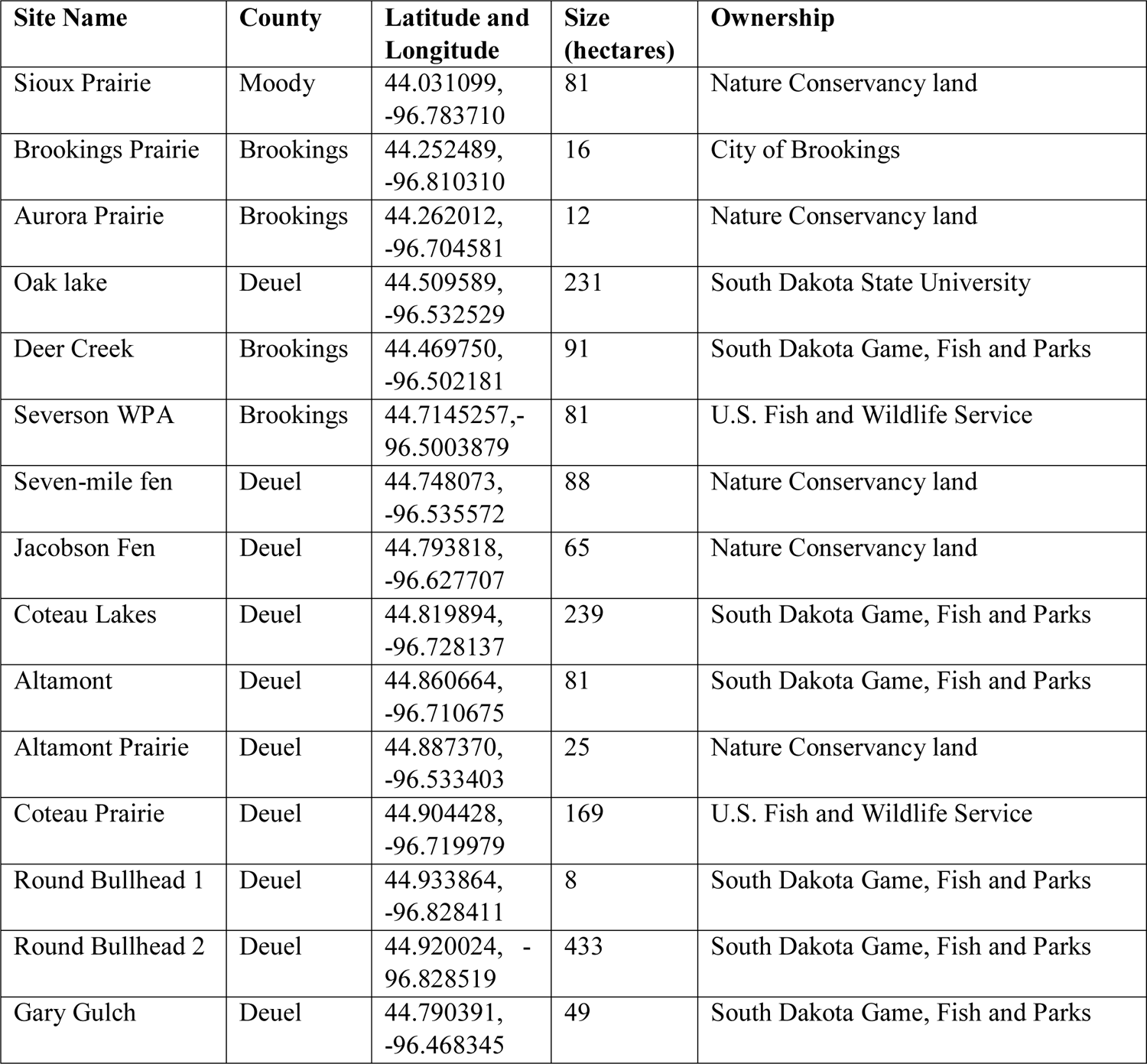
Description of temperate grassland sites in eastern South Dakota including site name, county name, latitude & longitude, size (in hectares) and ownership of sites.

### Data collection

#### Pollinator observation surveys

Between May and October of 2019, surveys were conducted for 30 minutes along 30 x 1 m transects on days warm enough to allow insect flight and in time periods when pollinators are expected to be active (15-35° C, between 08:00 – 17:00 hours). We divided the sampling into three seasons: early (May-June), mid (July-August) and late (September-October). Season intervals were selected based on consistent flowering phenology shifts found in the plant communities of the Prairie Coteau. For example, species belonging to the genera *Anemone (Ranunculaceae), Viola (Violaceae),* and *Sisyrinchium (Iridaceae)* predominately bloomed in the early season, while the mid and late seasons were dominated by species in the *Fabaceae* and *Asteraceae* (legume and sunflower families, respectively). Though these two families were predominately found in both seasons, the mid-season is distinct as this period marked a peak in the number of families in bloom with approximately six times more families present in our surveys in comparison to other seasons. Late season was characterized by a distinct shift in floral composition in which Asteraceae became the most prominent family in all sites with nearly all other families no longer flowering for the year. We sampled for one year and completed 114 transects.

From our roster of fifteen temperate grassland sites, we randomly sampled each site until flowering ceased at each location. Location and direction of transects were randomized at each visit using a list of randomly generated numbers to determine number of steps and cardinal directions before placing transects down. Transects were geospatial referenced using a Trimble Geo 7x GPS unit with 1-100 cm accuracy. We walked the entire length of the transect and recorded all plant-pollinator interactions from within one meter of the transect line on both sides. We defined pollinators as insect floral visitors that made contact with both the male and female reproductive parts of the flower, a commonly used criterion (Fenster et al. 2004). We documented each pollinator and the associated biotically-pollinated plant species when an interaction occurred. Additionally, we documented pollinator return visits to plants.

Pollinators were identified in situ to family and genus, then to morphospecies in order to quantify insect diversity. The pollinator observations in our study only focus on diurnal pollinators, however, this does not present a significant bias in our sampling. Our data set portrays a robust, representative sample of the plant-pollinator networks in this region considering only one species *(Silene vulgaris)* detected in our floral surveys (described in the next section) relies on nocturnal pollination, and this one species was only present in 1 transect of the 114 sampled. Insect voucher specimens were collected in the field with an aspirator and net, later identified to lowest taxonomic level and then categorized into functional groups. Specimens were identified using resources available through discoverlife.org, bugguide.net, and Key to the Genera of Nearctic Syrphidae (Miranda et al. 2013). Voucher specimens were verified for sampling completeness using the help of experts and the Severin-McDaniel Insect Research Collection available at South Dakota State University.

#### Floral Surveys

Floral surveys were conducted directly after insect pollinator observation surveys along the same transect with a 1 m^2^ quadrat. We placed the quadrat at each meter mark from 0 to 30 m and surveyed only one side of the transect considering each side mirrored the other with regards to species composition. Within the quadrat, we documented the presence of each biotically-pollinated plant species, number of individuals per species, number of flowering units defined as a unit of one (e.g., Ranunculaceae) or a blossom (e.g., Asteraceae) requiring a small pollinator to fly in order to access another flowering unit per species, and percent cover within quadrat per species. Plant voucher specimens were collected and identified using Van Bruggen (1985), verified with the help of experts (see *Acknowledgements),* and are curated at the C. A. Taylor Herbarium at South Dakota State University. Digitized plant collections for this study may be accessed on the Consortium of Northern Great Plains Herbaria (https://ngpherbaria.org/portal/).

To quantify the relationship between sugar resources at a site and pollinator diversity, nectar sugar content was collected using a handheld sucrose refractometer (Bellingham & Stanley Eclipse model) for each species during the growing season with a minimum of two samples per species. Nectar was collected in the field by collecting one flowering unit and compressing it into the refractometer lens to read volume and sucrose concentration based on the Brix scale. For composite flowers, the capitulum was first removed from the stem and sectioned into thirds. Then, a third of the ray and disc florets were compressed into the refractometer. Nectar volume was calibrated and standardized in lab using a micropipette and handheld sucrose refractometer. Volumes were categorized as low (<3.5 μl), medium (5.5 μl) and high (7.5 μl) based on how much liquid covered the main body lens of the refractometer during calibration. For example, a high liquid volume measuring at approximately 7.5 μl would completely cover the lens. These standardized volumes were applied to all biotically-pollinated plant species sampled in the field. Although these nectar measurements are a gross proxy of sucrose considering we are crushing the flowers to extract our measurements, they provided an efficient, consistent, and standardized method.

Nectar sugar was estimated by calculating an average nectar sugar concentration for each plant species and average volume from values recorded over the entire sampling season. We then estimated sugar for one flowering unit of each species by multiplying average volume and sugar concentration, and then converting to reflect grams of sugar per liter. Then, we calculated sugar for the entire plant for each species by multiplying estimated flower sugar and the number of flowers found on each transect. Finally, we summed all plant sugar found on each transect to obtain total transect sugar. Nectar data was not recorded for the following species: *Anticlea elegans, Helenium autumnale, Grindelia squarrosa, Lobelia spicata, Packera plattensis, Sisyrinchium montanum, Viola nephrophylla* as they were regionally rare and uncommon across sampling sites. When combined, all regionally rare species were found on approximately 9% of total transects. However, these species were accounted for in all other measurements implemented in the floral surveys listed in the previous section.

#### Pollinator and plant diversity

Pollinator and plant Shannon diversity were calculated using the ‘vegan’ package version 2.5-6 in R version 3.6.3 (R Core Team 2013; Oksanen et a. 2019). The Shannon index is calculated using the following formula:

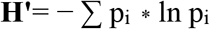

Where p_i_ is the proportions of each species found in a community and ln is the natural log. All diversity indices were natural log transformed. Shannon is the only diversity index presented in this study as it accounts for abundance, richness, and evenness. Pollinator diversity was calculated at the functional, family, genus, and species level by transect and season. Values used in pollinator diversity did not include return visits recorded during observation surveys. Likewise, plant diversity was calculated at the family, genus, and species level by transect and season using number of individuals recorded during floral surveys.

#### Landscape diversity and proportion

To estimate landscape diversity surrounding North American temperate grassland remnants, we used QGIS version 2.8.3 – with GRASS and ‘rgdal’, ‘maptools’, ‘rgeos’, ‘rasterVis’packages in R (Bivand and Rundel 2020a; Bivand et al. 2020b; Bivand and Lewin-Koh 2020c; Perpinan and Hijmans 2020). We obtained a 2019 cropland raster layer (CropScape; CDL) from USDA Ag Data Commons for South Dakota. We then created a vector data layer for the Prairie Coteau region from Google Earth Pro. The CDL 2019 raster layer was then clipped onto the Prairie Coteau layer and warped in order to view cropland use in our focus region with the correct projection (i.e., WGS 84/ UTM zone 14N). Further, we used QGIS to process and create a vector shapefile with each transect line collected from the Trimble Geo 7x.

We imported the vector shapefile produced in QGIS into R in order to create buffer layers at three different scales (500 m, 1000 m, and 3000 m) around each transect using the *gBuffer* function in the ‘rgeos’ package (Bivand and Rundel 2020a). The output of this function creates a table with pixel counts corresponding to each land-use in the CDL layer found within each transect buffer. The land-use pixel values were used to create Shannon diversity indices for each of our transects. Shannon diversity was calculated using the ‘vegan’ community ecology package version 2.5-6 in R. Landscape diversity was measured at the transect level due to the variation we found surrounding the transects in QGIS even at large scales (i.e., 1000 m and 3000 m).

We used the five most prominent land-uses (i.e., grassland, idle cropland, herbaceous wetlands, corn and soy) that comprise over 75% of the surrounding landscape to investigate the influence of land-use proportion on pollinator diversity. We used the same pixel values derived for landscape diversity to calculate land-use proportion for each transect. Land-use pixels were saved in Excel and proportion was calculated using simple equations for all scales (i.e., 500 m, 1000 m, 3000 m).

#### Network analysis

To quantify plant-pollinator community structure, we built quantitative visitation networks for each site by each of the three seasons. We collapsed all transects and created networks for each site by season considering that transect level analysis for network-level metrics does not provide enough interactions to include all sampled transects. By collapsing our samples into one season, we can include all observations and provide context of the entire community at a site in a given season in order to understand how network-level metrics are influenced by predictor variables (i.e., pollinator diversity) in our models. We focus on the mid-season in our results as it is the only season where all sites were able to be sampled more than once due to flooding conditions barring access to half of the sites in the early season. Sites were sampled only once during the late season, however, approximately half of the transects in the late season did not have enough interactions to derive network metrics. Nevertheless, we performed network analyses for the early and late seasons with halved sample sizes and found similar results to those found in the mid-season (Supplemental tables 1-3). The Deer Creek site was excluded from all network analyses as there were too few interactions even when transects were collapsed to generate network-level metrics.

Networks were constructed using a matrix of interactions between plants and pollinators including unique and return visits recorded during observation surveys. Documenting return visits allows us to quantify plant-pollinator communities using weighted network values that also account for visitation frequency. For each network, we calculated network specialization (H2’), connectance and nestedness. The H2’ specialization metric measures the degree of specialization at the community level (Blüthgen et al. 2006;2008) and is measured on a scale from 0 to 1, where a value approaching one indicates a network with increased selective interactions. Network-level specialization incorporates the interaction frequency between nodes and has been established as a more robust metric than species-level specialization given that network-level specialization is independent of network size (Blüthgen et al. 2006). Connectance represents the proportion of realized links out of all possible interactions within a network and is measured on a scale from 0 to 1. Connectance is used to describe the degree of generalization and interaction complexity within a network with a value approaching one indicating a more connected network with increased generalization (Dunne et al. 2002; Cusser and Goodell 2013). Nestedness describes a pattern of asymmetry in the interactions of a network (Bascompte et al. 2003).

Within a perfectly nested network, specialist species would interact with a subset of species that interact with generalists; thus, increasing the redundancy of interactions. Nestedness is measured on a scale from 0 to 100 with 0 describing perfect nestedness (Rodriguez-Girones & Santamaria 2006). Trends were analyzed using linear regressions. Using these network parameters, we can determine the influence of certain habitat attributes and the surrounding landscape on the preservation of specialized interactions and overall resilience of the community. All network metrics were calculated using the ‘bipartite’ package version 2.15 in R (Dormann et al. 2009).

## Statistical Analyses

### Relationship between landscape diversity, North American temperate grassland attributes and pollinator diversity

To quantify the relationship between pollinator diversity and attributes found within and outside of North American temperate grasslands, we used linear mixed effects models with site as the random effect. We used the packages ‘lme4’ version 1.1-23 and ‘nlme’ version 3.1-144 in R for the linear mixed effects models (Bates et al. 2015; Pinheiro et al. 2020). All variables in the model are at the transect level. The full model including both fixed and random effects was:

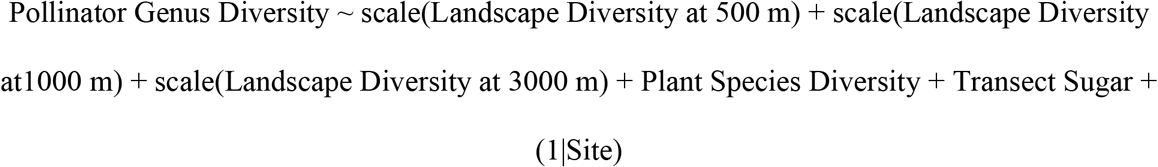

We then used stepwise-backward variable selection to simplify the model. The dropterm function in the ‘MASS’ package version 7.3-51.5 was used because it considers each variable individually and we can specify what test to use to compare the initial model as well as each of the possible alternative models with one less variable (Venables and Ripley 2020). For model assessment, we used AIC (Akaike Information Criterion) values and chi-square as criteria when dropping each variable one at a time from the full model stated above. Size and latitude were not included in the model to avoid collinearity. Thus, we used simple linear regressions for both size and latitude separately with pollinator diversity as the response variable.

### Relationship between landscape proportions, North American temperate grassland attributes and pollinator diversity

To quantify the relationship between landscape proportion, North American temperate grassland attributes and pollinator diversity, we used linear mixed effects models with site and season as the random effects. We used the packages ‘lme4’ and ‘nlme’ in R for the linear mixed effects models. All variables in the model are on the transect level. The full model including both fixed and random effects was:

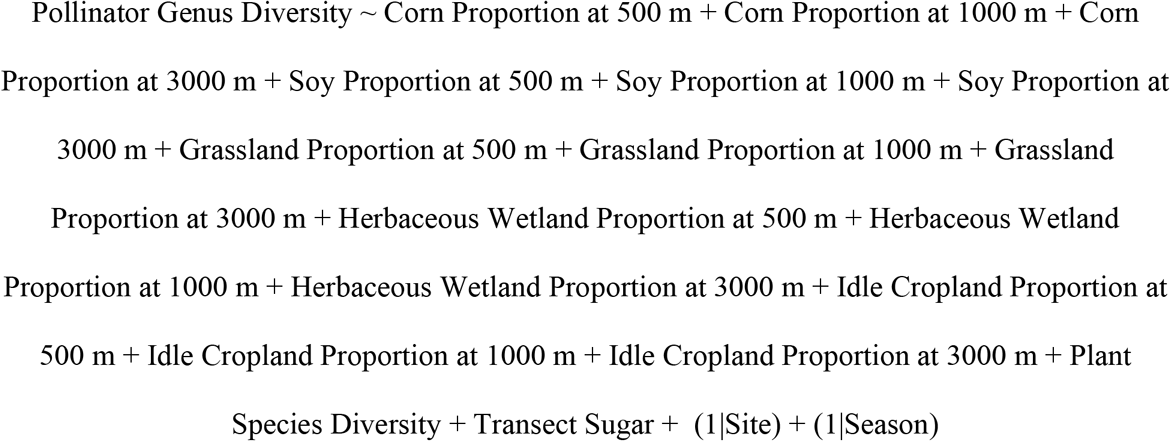

We implemented stepwise-backward variable selection to simplify this model. We incorporated the same model selection process that was applied to the landscape diversity model by using the dropterm function. For model assessment, we used AIC (Akaike Information Criterion) values and chi-square as criteria when dropping each variable one at a time from the full model stated above.

### Relationship between pollinator diversity and network structure

We used generalized linear models with a Gaussian distribution to determine the relationship between network structure and pollinator diversity due to some of our response variables being non-normally distributed. All variables were scaled to season level. Univariate regression models were used for each network parameter with pollinator diversity as the predictor variable and each network parameter as the response variable.

## Results

### North American temperate grassland attributes and the surrounding landscape

Landscape Shannon diversity varied from 0.47 to 2.10 across transects and spatial scales (Fig. 1). The most common land-use pixels found within the 3000 m buffers were grassland, herbaceous wetlands, fallow/idle cropland, corn, and soybean fields. When averaged across all sites at the 3000 m scale, grassland pixels covered 39%, fallow/idle cropland covered 11.5%, corn fields covered 9%, soybean fields covered 9%, and herbaceous wetlands covered 8% of 3000 m buffers. When combined, all five land-uses accounted for 76.5% of the landscape within 3000 m of our sites.

**Figure 1.**
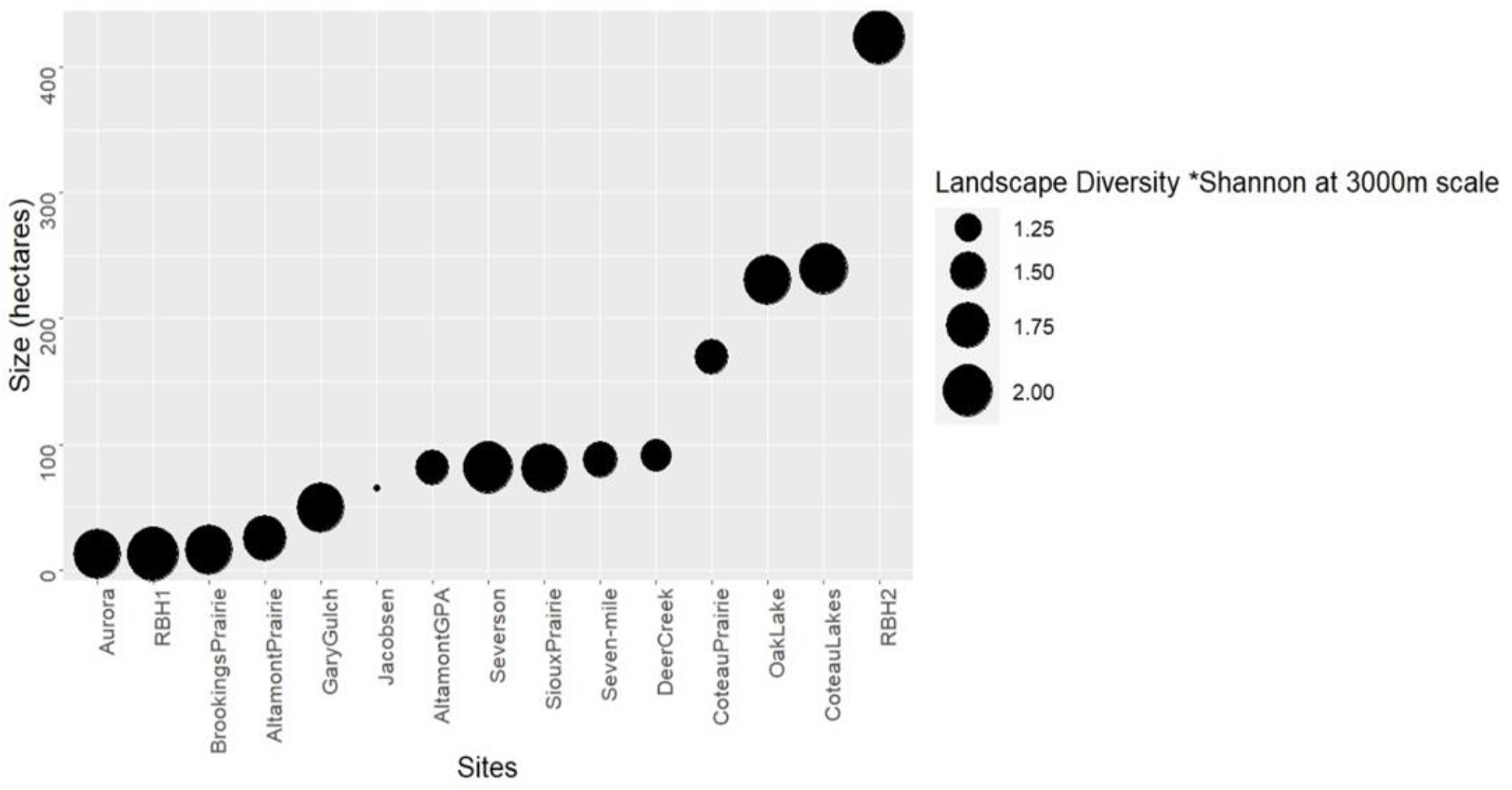
Size of remnant temperate grassland sites and the surrounding landscape diversity within 3000 meters of the sites in the Prairie Coteau Region near Brookings, South Dakota. Remnant temperate grasslands are ordered by size from smallest to largest in hectares. The size of the circle indicates the landscape diversity surrounding that site in 2019.

### Pollinator community

Among all 15 sites, 79 genera of pollinators representing 10 functional groups and 45 families were collected throughout the sampling season (see Appendix A for full list). 77% of total samples collected and observed were identified to genus. Samples that were not identified to genus were identified to the next taxonomic level (family 23%) and given a morphospecies classification which was used for pollinator genus analyses. Though we recorded morphospecies in the field, we found genus to be the lowest, most robust taxonomic level in the data set that could be identified with accuracy. Insect pollinators that could not be identified to genus were placed in a catch-all genus that consisted of the first five letters of their family name. For example, for a fly pollinator in the family Muscidae, we created a genus named Gen_Musci in the dataset in order to include these visitors in the analyses. We found pollinator genus diversity to be correlated with functional group diversity and family diversity (Supplemental Table 4). Thus, we focus on pollinator genus diversity in our analyses to reduce the number of analyses.

Shannon diversity of pollinator groups ranged from 0 to 2.2 among sites throughout all three seasons (Fig. 2). The majority of observations in the early season where diversity ranged from 0 – 1.7 included Syrphidae (36%), Muscidae (20%), Chloropidae (14.5%), and Halictidae (14%) (Supplemental Fig. 1). Mid-season diversity ranged from 0 – 2.2 with Apidae (27%), Syrphidae (25.7%), Cantharidae (12.5%), and Halictidae (11%) comprising the majority of observations (Supplemental Fig. 2). Late season diversity ranged from 0 – 1.8 with Syrphidae and Halictidae encompassing 58 and 28 percent of all observations, respectively (Supplemental Fig. 3).

**Figure 2.**
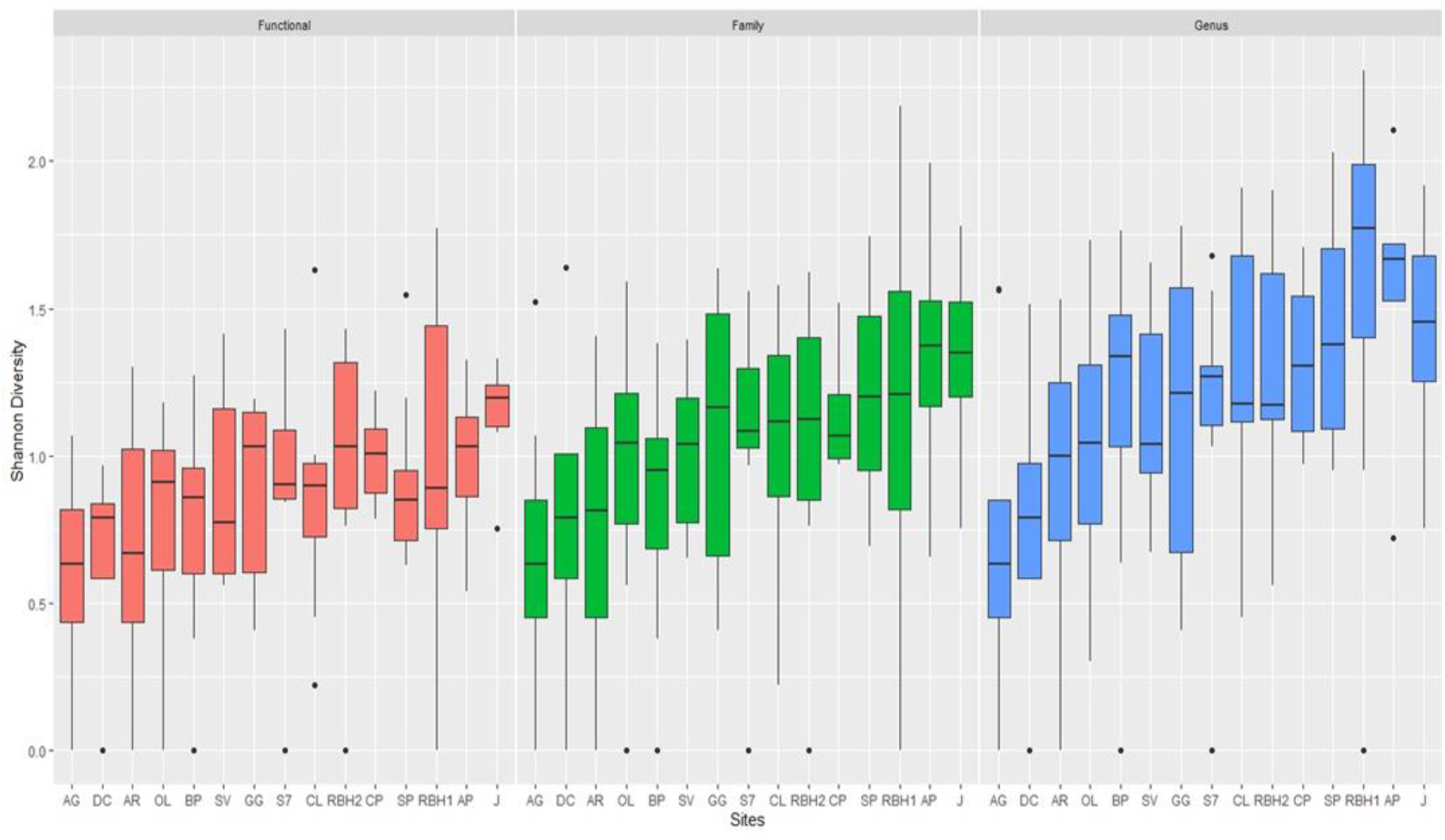
Shannon diversity of pollinators at the functional, family and genus level from May through October 2019 in the Prairie Coteau region near Brookings, South Dakota. Site abbreviations are as follows: DC = Deer Creek, AG = Altamont GPA, AR = Aurora, BP = Brookings Prairie, OL = Oak Lake Field Station, S7 = Seven-mile Fen, SV = Severson WMA, GG = Gary Gulch, CP = Coteau Prairie WMA, CL = Coteau Lakes GPA, RBH2 = Round Bullhead Lakes Site 2, SP = Sioux Prairie, AP = Altamont Prairie, RBH1 = Round Bullhead Site 1, J = Jacobsen Fen.R5T

### Plant community

Among all 15 sites, 87 species representing 24 families and 61 genera were collected throughout the growing season (see Appendix B for full list). Shannon diversity of biotically-pollinated plants ranged from 0 to 2 among sites throughout all three seasons (Fig. 3). Early season diversity ranged from 0 – 2 with *Anemone canadensis*, *Gallium boreali* and *Fragaria virginiana* as the most common species found on the transects (Supplemental Fig. 4). Mid-season diversity ranged from 0 – 2 with *Melilotus albus, Anemone canadensis,* and *Amorpha canescens* as the most common species found on the transects (Supplemental Fig. 5). Late season diversity ranged from 0 – 1.5 with *Symphyotrichum lanceolatum*, *Symphyotrichum ericoides* and *Heliopsis helianthoides* as the most common species found on the transects (Supplemental Fig. 6). We found that plant species diversity was correlated with family diversity and genus diversity (Supplemental Table 4). Thus, we focus on plant species diversity in our analyses to reduce the number of analyses.

**Figure 3.**
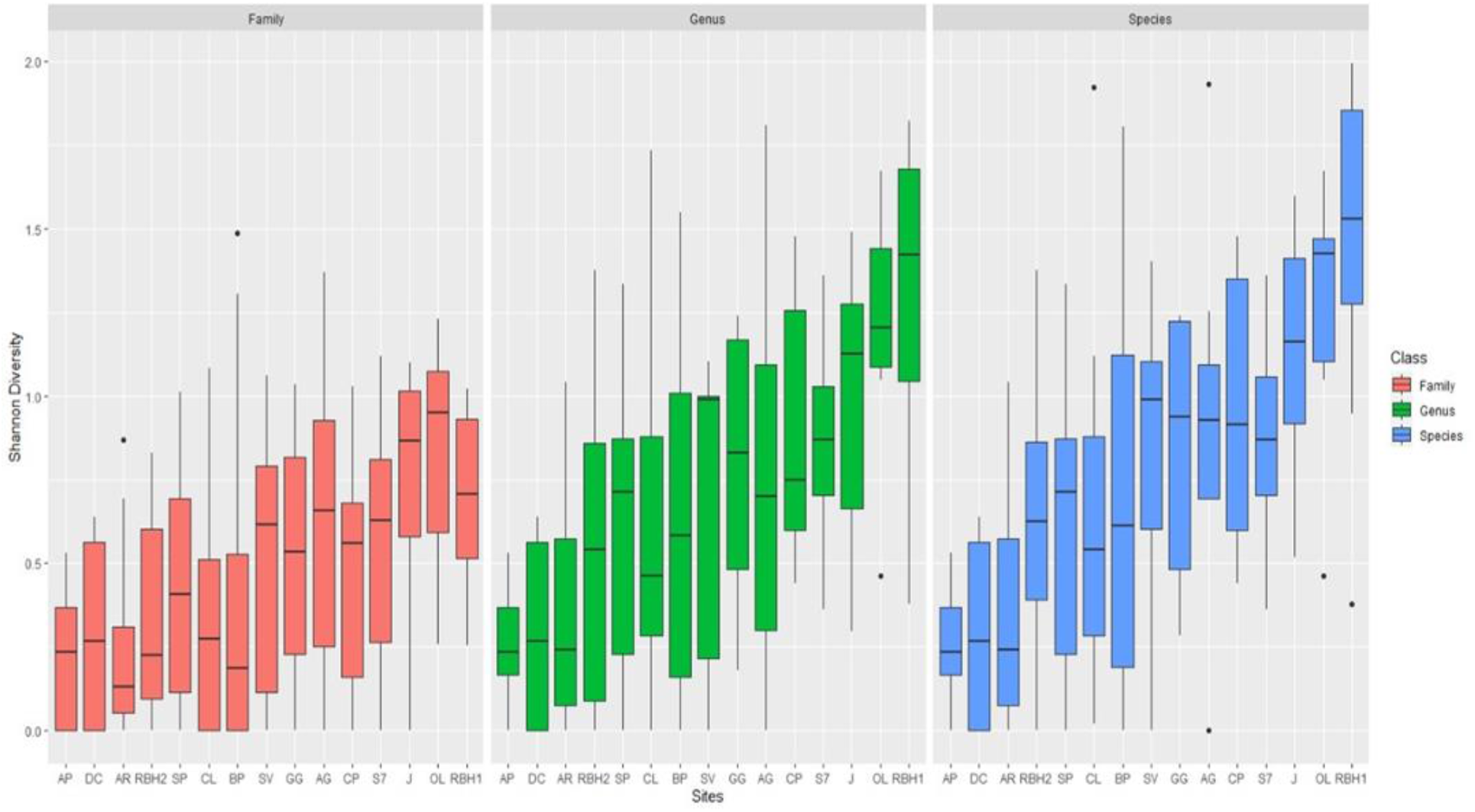
Shannon diversity of biotically-pollinated plants at the family, genus, and species level from May through October 2019 in the Prairie Coteau region near Brookings, South Dakota. Site abbreviations are as follows: DC = Deer Creek, AG = Altamont GPA, AR = Aurora, BP = Brookings Prairie, OL = Oak Lake Field Station, S7 = Seven-mile Fen, SV = Severson WMA, GG = Gary Gulch, CP = Coteau Prairie WMA, CL = Coteau Lakes GPA, RBH2 = Round Bullhead Lakes Site 2, SP = Sioux Prairie, AP = Altamont Prairie, RBH1 = Round Bullhead Site 1, J = Jacobsen Fen.

### North American temperate grassland attributes and landscape diversity effects on pollinator diversity

After model assessment, we found the final, simplified model explaining pollinator genus diversity included transect sugar as the only predictor variable (Table 2). Transect sugar demonstrates a significant positive relationship with pollinator genus diversity (Table 3). We found no relationship between pollinator diversity and North American temperate grassland size or latitude.

**Table 2.** North American temperate grassland attributes and landscape diversity effects on pollinator diversity in the Prairie Coteau region near Brookings, South Dakota in 2019. Pollinator genus diversity is the response variable in the following models and refers to the Shannon diversity of pollinators in the sampled remnant temperate grasslands within the Prairie Coteau region. The landscape diversity variables refer to the scales at which we measured landscape diversity (500 m, 1000 m, 3000 m). Plant species diversity refers to the Shannon diversity of biotically-pollinated plant species in the sampled remnant temperate grasslands and Transect Sugar refers to the amount of sugar that was available on each transect. Transect sugar values were derived from the nectar estimations collected from floral surveys. Random effects in the model include the remnant temperate grassland site itself. Backwards stepwise selection is applied to the following models in the table using single term deletion in the dropterm function in ‘MASS’ package in R. We list each model in the table and its associated AIC value. The first AIC values refer to the AIC score of the current model listed. The AIC values next to each variable refer to the AIC score of the model if the associated variable was removed. The p-values in the table are outputs from the likelihood ratio test for each variable. * values indicate that removing the associated variable would significantly change the model. *lndicates significance at the α = 0.05 level. P < 0.05; * P < 0.05; ** P < 0.01; *** P < 0.001

| **Model Selection for Pollinator Genus Diversity =** | **Likelihood ratio test** | **AIC** | **p-value** |
| --- | --- | --- | --- |
| **Landscape Diversity at 500 m + Landscape Diversity at 1000 m + Landscape Diversity at 3000 m + Plant Species Diversity + Transect Sugar** |  | 177.79 |  |
| Landscape Diversity at 500 m | 1.32 | 177.12 | 0.250 |
| Landscape Diversity at 1000 m | 0.28 | 176.07 | 0.596 |
| Landscape Diversity at 3000 m | 0.02 | 175.81 | 0.881 |
| Plant Species Diversity | 0.68 | 176.47 | 0.409 |
| Transect Sugar | 4.43 | 180.22 | 0.035* |
| **Landscape Diversity at 500 m + Landscape Diversity at 1000 m + Plant Species Diversity + Transect Sugar** |  | 175.81 |  |
| Landscape Diversity at 500 m | 1.35 | 175.16 | 0.246 |
| Landscape Diversity at 1000 m | 0.41 | 174.22 | 0.522 |
| Plant Species Diversity | 0.66 | 174.48 | 0.415 |
| Transect Sugar | 4.42 | 178.23 | 0.036* |
| **Landscape Diversity at 500 m + Plant Species Diversity + Transect Sugar** |  | 174.22 |  |
| Landscape Diversity at 500 m | 1.02 | 173.24 | 0.313 |
| Plant Species Diversity | 0.72 | 172.94 | 0.396 |
| Transect Sugar | 4.30 | 176.53 | 0.038* |
| **Landscape Diversity at 500 m + Transect Sugar** |  | 172.94 |  |
| Landscape Diversity at 500 m | 1.01 | 171.95 | 0.315 |
| Transect Sugar | 4.61 | 175.56 | 0.032* |
| **Transect Sugar** |  | 171.95 |  |
| Transect Sugar | 6.41 | 176.37 | 0.011* |

**Table 3.** Linear mixed effect analysis of transect sugar on pollinator genus Shannon diversity in the Prairie Coteau region near Brookings, South Dakota. P < 0.05; * P < 0.05; ** P < 0.01; *** P < 0.001

| Variables | Coefficient | Standard Error | Degrees of Freedom | p-value |
| --- | --- | --- | --- | --- |
| Intercept | 1.16 | 0.06 | 14.78 | 5.97e-12*** |
| Transect Sugar | 0.03 | 0.01 | 104.86 | 0.013* |

### North American temperate grassland attributes and land-use proportion effects on pollinator diversity

We found the final, simplified model explaining pollinator genus diversity included corn proportion at 500 m, soy proportion at 1000 m, herbaceous wetland proportion at 1000 m, and transect sugar (Table 4). All predictor variables in the model demonstrated a positive relationship with pollinator genus diversity (Table 5).

**Table 4.** North American temperate grassland attributes and landscape proportion effects on pollinator diversity in the Prairie Coteau region near Brookings, South Dakota in 2019. Pollinator genus diversity is the response variable in the following models and refers to the Shannon diversity of pollinators in the sampled remnant temperate grasslands within the Prairie Coteau region. Landscape proportion for the five most prominent land-uses were measured for the same scales used for landscape diversity (500 m, 1000 m, 3000 m). Plant species diversity refers to the Shannon diversity of biotically-pollinated plant species in the sampled remnant temperate grasslands and Transect Sugar refers to the amount of sugar that was available on each transect. Transect sugar values were derived from the nectar estimations collected from floral surveys. Random effects in the model include season (i.e., early, mid and late) and the remnant temperate grassland site itself. Backwards stepwise selection was applied to the initial model presented in the table using single term deletion in the dropterm function in ‘MASS’ package in R. We present the initial model in the table and then the final, simplified model after single term deletion. The AIC values at the top refer to the AIC score of the current model listed. The AIC values next to each variable refer to the AIC score of the model if the associated variable was removed. The p-values in the table are outputs from the likelihood ratio test for each variable. * values indicate that removing the associated variable would significantly change the model. *lndicates significance at the α = 0.05 level. P < 0.05; * P < 0.05; ** P < 0.01; *** P < 0.001

| **Model Selection for Pollinator Genus Diversity =** | **Likelihood**<br>**ratio test** | **AIC** | **p-value** |
| --- | --- | --- | --- |
| Initial Model:<br>**Corn Proportion at 500 m + Corn Proportion at 1000 m +**<br>**Corn Proportion at 3000 m + Soy Proportion at 500 m + Soy**<br>**Proportion at 1000 m + Soy Proportion at 3000 m +**<br>**Fallow/Idle Cropland Proportion at 500 m + Fallow/Idle**<br>**Cropland Proportion at 1000 m + Fallow/Idle Cropland**<br>**Proportion at 3000 m + Grassland Proportion at 500 m +**<br>**Grassland Proportion at 1000 m + Grassland Proportion at**<br>**3000 m + Herbaceous Wetland Proportion at 500 m +**<br>**Herbaceous Wetland Proportion at 1000 m + Herbaceous**<br>**Wetland Proportion at 3000 m + Plant Species Diversity +**<br>**Transect Sugar** |  | 185.90 |  |
|  | 1.71 | 185.61 | 0.19 |
| Corn Proportion at 500 m | 0.67 | 184.57 | 0.41 |
| Corn Proportion at 1000 m | 0.30 | 184.19 | 0.59 |
| Corn Proportion at 3000 m | 0.03 | 183.93 | 0.86 |
| Fallow/Idle Cropland Proportion at 500 m | 0.17 | 184.07 | 0.68 |
| Fallow/Idle Cropland Proportion at 1000 m | 3.22 | 187.12 | 0.07 |
| Fallow/Idle Cropland Proportion at 3000 m | 0.48 | 184.38 | 0.49 |
| Grassland Proportion at 500 m | 0.32 | 184.22 | 0.57 |
| Grassland Proportion at 1000 m | 0.58 | 184.48 | 0.45 |
| Grassland Proportion at 3000 m | 0.76 | 184.66 | 0.38 |
| Herbaceous Wetland Proportion at 500 m | 2.52 | 186.42 | 0.11 |
| Herbaceous Wetland Proportion at 1000 m | 0.60 | 184.50 | 0.44 |
| Herbaceous Wetland Proportion at 3000 m | 0.09 | 183.99 | 0.76 |
| Soy Proportion at 500 m | 2.61 | 186.51 | 0.11 |
| Soy Proportion at 1000 m | 0.68 | 184.58 | 0.41 |
| Soy Proportion at 3000 m | 1.16 | 185.06 | 0.28 |
| Plant Species Diversity | 6.04 | 189.94 | 0.01 * |
| Transect Sugar |  |  |  |
| Final Model:<br>**Corn Proportion at 500 m + Herbaceous Wetland Proportion**<br>**at 1000 m + Soy Proportion at 1000 m + Transect Sugar** |  | 167.22 |  |
| Corn Proportion at 500 m | 6.90 | 172.12 | 0.009 ** |
| Herbaceous Wetland Proportion at 1000 m | 4.92 | 170.14 | 0.026 * |
| Soy Proportion at 1000 m | 6.82 | 172.04 | 0.009 ** |
| Transect Sugar | 5.99 | 171.20 | 0.014 * |

**Table 5.**
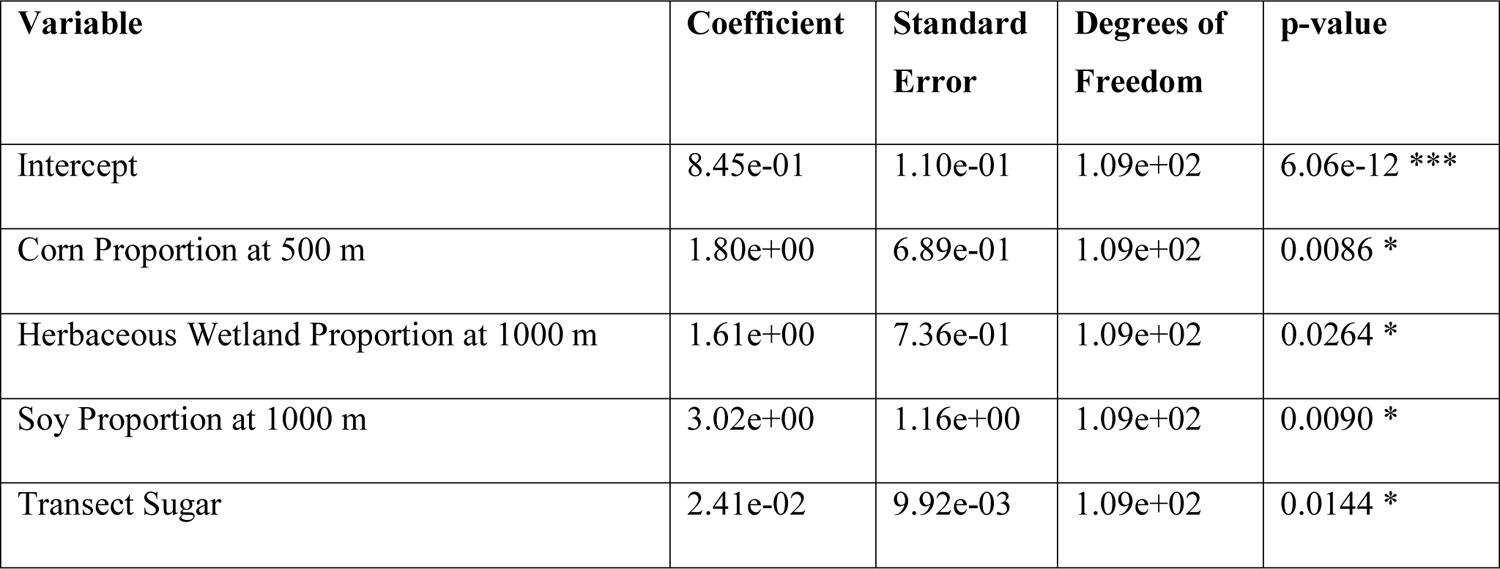
Linear mixed effects analysis results of land-use proportions and transect sugar on pollinator genus Shannon diversity in the Prairie Coteau region near Brookings, South Dakota. P < 0.05; * P < 0.05; ** P < 0.01; *** P < 0.001

| Variable | Coefficient | Standard Error | Degrees of Freedom | p-value |
| --- | --- | --- | --- | --- |
| Intercept | 8.45e-01 | 1.10e-01 | 1.09e+02 | 6.06e-12 *** |
| Corn Proportion at 500 m | 1.80e+00 | 6.89e-01 | 1.09e+02 | 0.0086 * |
| Herbaceous Wetland Proportion at 1000 m | 1.61e+00 | 7.36e-01 | 1.09e+02 | 0.0264 * |
| Soy Proportion at 1000 m | 3.02e+00 | 1.16e+00 | 1.09e+02 | 0.0090 * |
| Transect Sugar | 2.41e-02 | 9.92e-03 | 1.09e+02 | 0.0144 * |

### Plant-pollinator network analysis

From May through October, we observed 518 unique plant-pollinator interactions and a total of 9,877 observations of pollinators visiting plants. The most common floral visitors throughout the entire sampling period were Syrphidae (31%), Apidae (19%), and Halictidae (13%). The plant species with the most interactions throughout the entire sampling period include *Melilotus albus* (20.7%), *Amorpha canescens* (10.6 %), *Euphorbia esula* (8.5%), and *Achillea millefolium* (6.7%).

### Network metrics and pollinator genus diversity

Network-level specialization (H2’) ranged from 0.35 – 0.81 between temperate grassland sites during mid-season. Pollinator genus diversity had a significant positive relationship with network-level specialization (p < 0.002, t = 4.313, slope coefficient: 0.304 ± 0.072) (Fig. 4). Connectance ranged from 0.10 – 0.29 between temperate grassland sites at the mid-season. Pollinator genus diversity demonstrated a negative relationship with connectance (p < 0.04, t = – 2.39, slope coefficient: −0.077 ± 0.032) (Fig. 4). Nestedness ranged from 9 – 25 between temperate grassland sites at the mid-season. We found no significant relationship between pollinator genus diversity and nestedness (p < 0.50, t = −0.70, slope coefficient: −3.015 ± 4.29).

**Figure 4.**
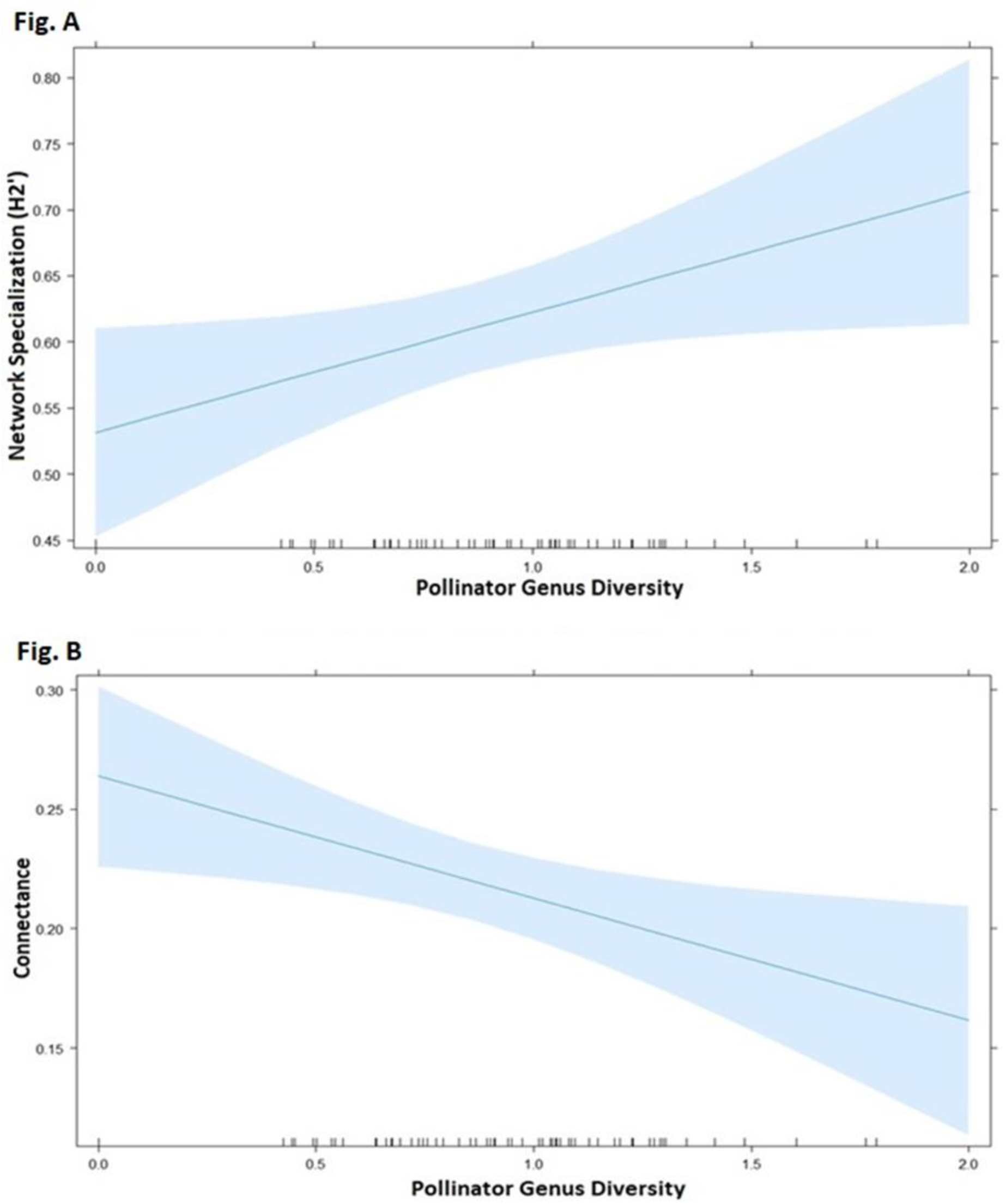
(A) Effect of pollinator genus diversity on network-level specialization (H2’) in the plant-pollinator communities found within the Prairie Coteau region near Brookings, South Dakota in 2019. Pollinator genus diversity refers to the Shannon diversity of pollinators in the sampled remnant temperate grasslands within the Prairie Coteau region. Network specialization ranges from 0 −1 with values approaching 1 denoting more specialized interactions in a community. Pollinator genus diversity has a significant positive effect on network-level specialization at the mid-season (p < 0.002, t = 4.313, std. error = 0.07). (B) Effect of pollinator genus diversity on connectance in the plant-pollinator communities found within the Prairie Coteau region near Brookings, South Dakota in 2019. Pollinator genus diversity refers to the Shannon diversity of pollinators in the sampled remnant temperate grasslands within the Prairie Coteau region. Connectance ranges from 0 – 1 with the value of 1 indicating a maximally connected community. Pollinator genus diversity has a significant negative effect on network connectance (p < 0.04, t = −2.39, std. error = 0.032).

## Discussion

Our study expands on previous research (Aizen et al. 2003; Ashworth et al. 2004; Heard et al. 2007; Carvalheiro et al. 2010; Ferreira et al. 2013; Boyle et al. 2019; Ponisio et al. 2019; Twerd and Banaszak-Cibicka 2019) documenting the influence of landscape modification and habitat fragmentation on the persistence of plant-pollinator communities in natural habitats through changes in pollinator availability and diversity. We further develop these approaches by investigating a combination of both local habitat attributes (i.e., patch size, latitude, plant species diversity, nectar resources) and landscape effects on pollinator diversity and linking the effects of pollinator diversity to the structure of plant-pollinator communities using a network-based approach. Our results indicate the most important habitat attribute predicting pollinator genus diversity in the Prairie Coteau region is the amount of floral sugar found at remnant temperate grassland sites as extrapolated from transect sampling. Additionally, we found landscape diversity had no effect on pollinator genus diversity. Instead, the proportion of certain land-uses surrounding remnant temperate grassland sites had a significant effect on pollinator genus diversity. In turn, pollinator genus diversity demonstrated a positive effect for overall network specialization and consequentially a negative effect on connectance. Below, we discuss the local and landscape effects on pollinator diversity followed by the effects of pollinator diversity on overall community structure and how our findings may influence future planning for the conservation of temperate grassland remnants and pollinator diversity in the Northern Great Plains.

### The role of local resources and North American temperate grassland attributes on pollinator diversity

Nectar resources (i.e., plant sugar) were an important determinant of pollinator diversity at remnant temperate grassland sites in the agriculturally dominated landscape of the Northern Great Plains. These findings indicate floral abundance and nectar rewards influence pollinator behavior, also observed by others (Potts et al. 2009; Fowler et al. 2016), especially in disturbed, fragmented habitats (Potts et al. 2003; Stang et al. 2006; Ouvrard et al. 2018). The common, non-native forb *Melilotus albus* recorded in 10 of our 15 sites, had greater average nectar contribution per transect (based on number of flowers and sugar concentration per flower) and comprised 20% of total interactions throughout the entire sampling season, making it a core node in our plant-pollinator networks. Our findings with *M. albus* are also consistent with other studies demonstrating bees will heavily utilize non-native plants in disturbed habitats, even though they do not exhibit a preference for non-native plants over native plants (Hinners et al. 2009; Williams et al. 2011). Considering the impacts non-native plant species can have on pollinators, it is imperative we consider the role that non-native plants play in shaping community interactions, particularly in heavily disturbed and modified landscapes such as those dominated by agriculture. The role of non-native pollinator species in plant-pollinator networks should also be examined considering honey bees could compete with native pollinators for foraging resources (Sugden et al. 1996; Thomson 2004). In the Northern Great Plains, a study conducted from 2015-2017 found both honey bees and wild bees visiting *Melilotus* with honey bees utilizing *Melilotus* more than wild bees (Otto et al. 2020). However, honey bees in our study were only present in 7 out of 15 temperate grassland sites and made up 12% of total interactions. Honey bee observations only occurred due to presence of commercial honey bee hives within 1 km of our sites and overall, account for a low frequency of visitation in networks.

*Melilotus* is considered an attractive resource to insect pollinators because of their high nectar production, strong scent, and abundance of flowers produced with up to 350,000 flowers per plant (Royer and Dickinson 1999; Spellman et al. 2015). As *Melilotus* continues to establish within the Northern Great Plains, we can expect it to become a prominent, stabilizing anchor in plant-pollinator networks that could facilitate increased pollinator visitation and diversity (Tepedino and Griswold 2008; Spellman et al. 2015). Other invasive species in temperate grasslands such as *Cirsium arvense* have been found to influence the topology of plant-pollinator networks by decreasing modularity, which is associated with greater network stability; however, the effects of non-native plant species on native plant reproduction were not investigated (Larson et al. 2016). Despite the warranted concern over the impacts of invasive plant species on native plant communities (i.e., reduction of pollen quality and quantity) (Waser 1983; Campbell 1985; Brown et al. 2002; Morales and Traveset 2008; Kandori et al. 2009), some studies have shown that *Melilotus* can benefit community interactions by increasing pollinator visitation, diversity, and abundance (Morales and Traveset 2005; Tepedino and Griswold 2008; Spellman et al. 2015). In the Northern Great Plains, Otto et al. (2020) found both species of *Melilotus* were among the three most abundant flowering species in their transects, corresponding with our findings that *Melilotus* is a core node in the plant-pollinator networks in our study region.

### Landscape effects on pollinator diversity

Landscape diversity did not exhibit a relationship with pollinator genus diversity at any scale (i.e., 500 m, 1000 m, and 3000 m). However, we found that increased proportions of corn fields at 500 m, soybean fields at 1000 m, and herbaceous wetlands at 1000 m demonstrated a positive effect on pollinator genus diversity. Herbaceous wetlands adjacent to perennial grasslands, specifically those found in the Upper Midwest and Prairie Coteau regions, are valuable resources for insect pollinators and help increase native bee diversity and abundance (Vickruck et al. 2019; Begosh et al. 2020). The positive relationship between pollinator genus diversity and proportion of herbaceous wetlands in the Prairie Coteau may be related to the resources provided by the semi-natural habitats within the landscape.

Conversely, intensive agricultural fields (i.e., corn and soy fields) do not provide ample foraging or nesting resources for pollinators throughout the growing season and have been documented to negatively affect the hive health of certain pollinators (e.g., bumblebees) by reducing the quality and diversity of pollen available (Hass et al. 2018). The rise in pollinator genus diversity as intensive agriculture increases at various scales (i.e., 500 m and 1000 m) in our study could indicate an oasis effect occurring where insect pollinators are crossing substantial distances to forage on the ample nectar resources provided by temperate grassland patches in the Northern Great Plains (Bock et al. 2008; Gotlieb et al. 2011). Approximately 50% of the interactions in our plant-pollinator networks were dominated by syrphid flies and apid bees, which are highly mobile and utilize a variety of habitat types, indicating the insect pollinators in our observed networks may have traveled considerable distances to utilize the semi-natural reservoirs embedded within this cropland matrix. (Morris 1993; Sommaggio et al. 1999; Beekman and Ratneiks 2000; Klecka et al. 2018; Jauker et al. 2019). These findings are also consistent with other studies such as Winfree et al. (2007) and Hoehn et al. (2008) demonstrating that landscapes dominated by intensive agriculture can still maintain relatively diverse pollinator communities provided patches of floral resources were present.

### How does pollinator diversity influence the overall plant-pollinator community structure?

While other studies have focused on the botanical side of plant-pollinator networks in order to investigate network structure (Larson et al. 2016; Robinson et al. 2018; Cunningham-Minnick and Crist 2020; Hernandez-Castellano et al. 2020), we instead use pollinator diversity as our predictor variable to investigate network structure in perennial grasslands in the Northern Great Plains. The availability and abundance of nectar resources influence pollinator behavior which can potentially impact community structure on a larger scale. For example, Nottebrock et al. (2017) found that pollinator visitation rates strongly depended on phenological variation of site□scale sugar amounts. Further, the seed sets of their focal plants increased with both site-scale sugar amounts and nectar sugar amounts of conspecific neighbors. While their study did not investigate network structure, it demonstrates that the quantity and availability of nectar resources can have broader community impacts for both plants and pollinators. Our results indicate pollinator genus diversity influences connectance and network specialization (H2’) in reinforcing ways, with connectance decreasing and specialization increasing with greater pollinator genus diversity. We did not observe any relationship between nestedness and pollinator genus diversity, however we did find that networks were nested, as expected of asymmetric, mutualistic networks which contain hubs (i.e., highly linked species) that form a central core of interactions linking other species within the network (Bascompote et al. 2003; Jordano et al. 2003). Networks in our study fell within a range which indicates a nested pattern (9-25) that has also been observed by others within working grasslands (Vanbergen et al. 2014; Jauker et al. 2019).

Our findings align with predictions by Montoya and Yvon-Durocher (2007) who suggested that interspecific competition among pollinators may be driving network specialization more so than floral diversity (see also Fründ et al. 2010; Valdovinos et al. 2016). Network-level specialization can be beneficial for plant-pollinator communities as it may allow for increased efficiency of pollen transfer which can reflect a higher resilience and stability in the community (Arceo-Gomez 2020). Likewise, the negative relationship between connectance and pollinator genus diversity suggests plant-pollinator networks in the Prairie Coteau region can be prone to being less complex and connected (Dunne et al. 2002). Nested networks displaying a higher degree of connectance are considered more resilient and stable, making them important considerations for conservation value (Memmott et al. 2004; Okuyama and Holland 2008; Thébault and Fontaine 2010). The nested pattern found in the networks indicates a degree of interaction redundancy that likely contributes to community stability (Bascompte et al. 2003; Nielsen and Bascompte 2007). Nonetheless, nestedness does not seem to be influenced by the diversity of pollinators in these remnant temperate grassland communities. Overall, our results suggest that temperate grasslands with increased pollinator diversity can maintain more specialized plant-pollinator interactions that can potentially promote resilient plant-pollinator communities in the Prairie Coteau region.

## Conservation implications

Our results demonstrate that managing pollinator resources (sucrose availability) within remaining habitats and the composition of the surrounding landscape should be carefully considered when seeking to promote pollinator diversity and preserve pollination services. These results agree with other studies (Potts et al. 2005; Tscharntke et al. 2005; Potts et al. 2009; Heard et al. 2007; Cusser and Goodell 2013; Moreira et al. 2015; Smart et al. 2016), which identified how pollinator populations are influenced by a series of interacting factors (i.e., landscape configuration and floral community composition), rather than a singular attribute such as patch size. Landscape conversion in agricultural regions is not likely to diminish soon, making it imperative that we recognize the conservation value of the remaining habitat fragments such as those found in the Prairie Coteau.

Our study demonstrated plant species with high nectar rewards promoted pollinator diversity within temperate grasslands of the Northern Great Plains. While we cannot conclude whether non-native nectar resources provide more benefits to the overall health of pollinators in this region, future research should address the effects of native vs. non-native floral resources on pollinator health and network community structure in disturbed, fragmented habitats. Several studies (Lopezaraiza-Mikel et al. 2007; Aizen et al. 2008; Bartomeus and Santamaria 2008; Padron et al. 2009) have indicated that non-native plants may alter the structure of plant-pollinator networks by monopolizing the interactions of the most generalist pollinators. Recently, Hernandez-Castellano et al. (2020) and Cunningham-Minnick and Crist (2020) found that the integration of non-native plant species into plant-pollinator networks could dilute pollination services for native plants, with potential risks for pollinators that may not be able to use invasive floral resources. Thus, it would be valuable to apply this question to long-term monitoring of non-native plant species and their pollinators to determine if there will be any consequential changes to network structure over time. Long-term network studies are not common (Burkle and Alarcón 2011) and the matter of network stability in agriculture-dominated landscapes should be further investigated to ascertain how local and landscape changes affect plant-pollinator networks on a wider temporal scale.

While current evidence indicates network metrics remain stable across years (Alarcón et al. 2008; Petanidou et al. 2008; Burkle and Irwin 2009; Dupont et al. 2009b), consideration should be made for temporal shifts in plant and pollinator communities that could occur due to climate change (Roy and Sparks 2000; Fitter and Fitter 2002; Forister and Shapiro 2003; Bartomeus et al. 2011; Calinger et al. 2013; CaraDonna et al. 2014). Changes in flowering time and pollinator activity could result in temporal mismatching between actors in a network, potentially shifting network structure and stability (Albrecht et al. 2010; Morton and Rafferty 2017). Network interactions are considered fairly dynamic. Overall network structure has shown to remain stable even when network ‘rewiring’ occurs as a result of shifts in actors due to changes in abundance of plants or pollinators between years (Petanidou et al. 2008; Dupont et al 2009b); however, most multi-year network studies only extend to 2-4 years (Burkle and Alarcón 2011). As ecosystems continue to experience habitat disturbance on various scales along with climate change, it would be valuable to understand if long-term changes to species compositions would impact the structure of future plant-pollinator networks. Additionally, further research on understanding the influence of land management strategies (e.g., prescribed burning and grazing) within remnant, semi-natural habitats could provide insight on resource changes and availability at a local scale.

Native pollinators provide valuable ecosystem services for both native plant communities and agricultural fields by increasing plant reproductive success and crop yield (Fontaine et al. 2005; Albrecht et al. 2007; Albrecht et al. 2012; Rogers et al. 2014). Increasing pollinator abundance and diversity while maintaining specialized interactions can promote stability of this essential ecosystem service by enhancing the overall visitation and pollination of the plant communities present (Klein et al. 2007; Hoehn et al. 2008; Albrecht et al. 2012). Conserving these pollination services is important for sustaining future food security of the 35% of agricultural crops that rely on pollination globally, in addition to local crops such as sunflower and soybean which are economically valuable for the Northern Great Plains region (Klein et al. 2007; Vanbergen et al. 2013; Mallinger et al. 2019). Additionally, the protection and maintenance of ecosystem services (Costanza et al. 1998) directly supports ecosystem function (Peterson et al. 2010). Land managers should bear in mind the influence of nectar rewards on pollinator movement and behavior, and develop decisions focused on providing ample resources within the landscape. Furthermore, the conservation of other semi-natural habitats (i.e., herbaceous wetlands), which provide resources beyond nectar rewards should be prioritized to promote diverse pollinator communities, at least in the Northern Great Plains. The oasis effect demonstrated in our study will likely only intensify as time passes and the role of habitat reservoirs, like the temperate grassland remnants, will become increasingly important as oases for pollinator communities in agriculture-dominated regions.

## ACKNOWLEDGMENTS

SDSU and the Prairie Coteau region is located on the ancestral territory of the Oceti Sakowin, commonly called Sioux. We thank S. Stiles, J. Gelderman, T. Wallner, N. Petersen, E. Baier, and S. Daniels for their field assistance, J. Purntun for providing botanical expertise, landowners and managers that provided permissions and advice on site selection. Funding of this project was by several Hatch grants and the North Central Sun Grant Initiative (USDA/DOE) SA1500640.

## Appendices

## Appendix A

The table below lists all of the pollinators observed and identified in the Prairie Coteau region near Brookings, South Dakota in 2019 down to lowest taxonomic level. Insect pollinators that could not be identified to genus were placed in a catch-all genus that consisted of the first five letters of their family name. For example, for a fly pollinator in the family Muscidae, we created a genus named Gen_Musci in the dataset in order to include these visitors in the analyses. Table also includes number of remnant temperate grassland sites the pollinators were present and the number of total observed interactions of each pollinator from May through October 2019.

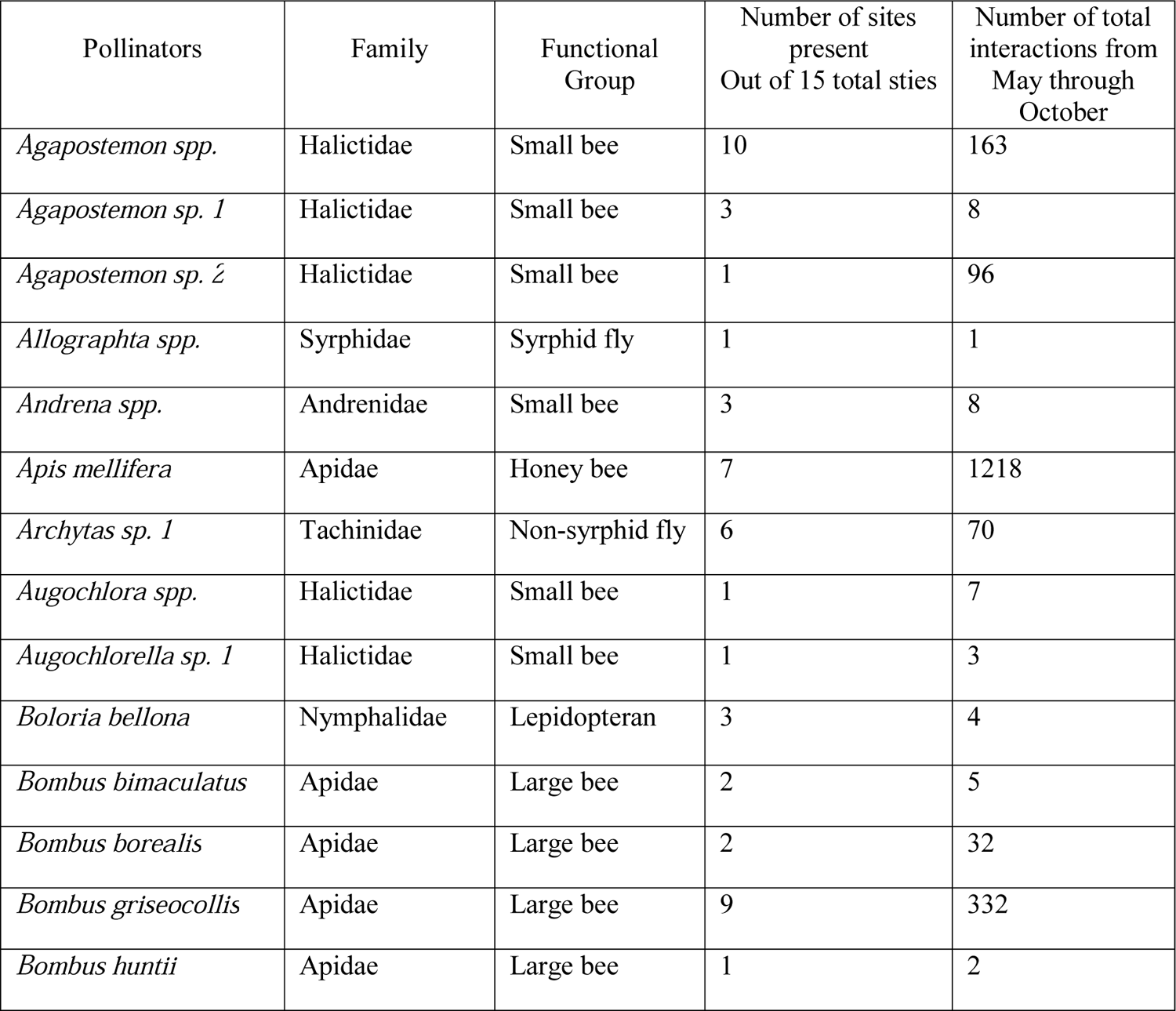
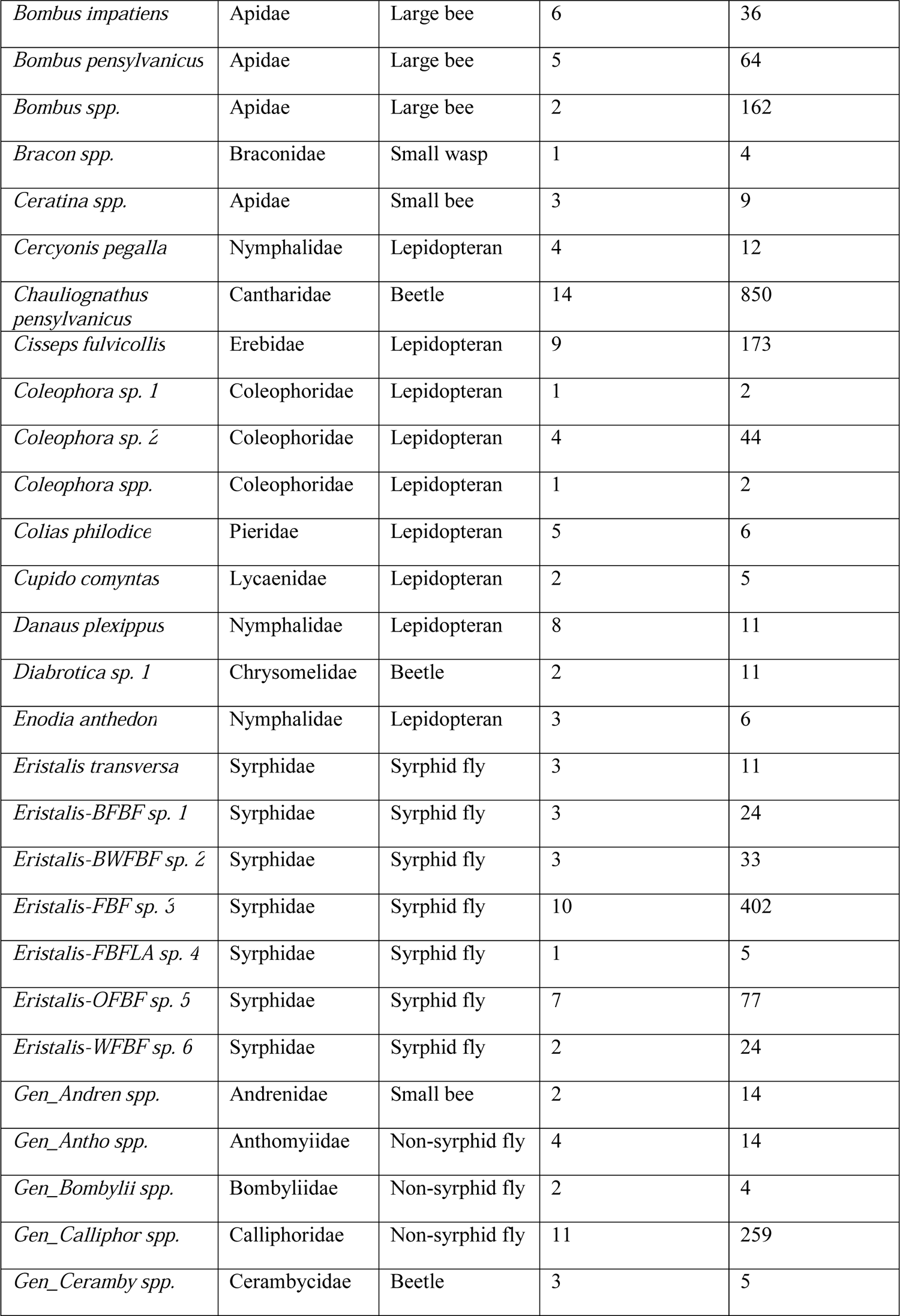
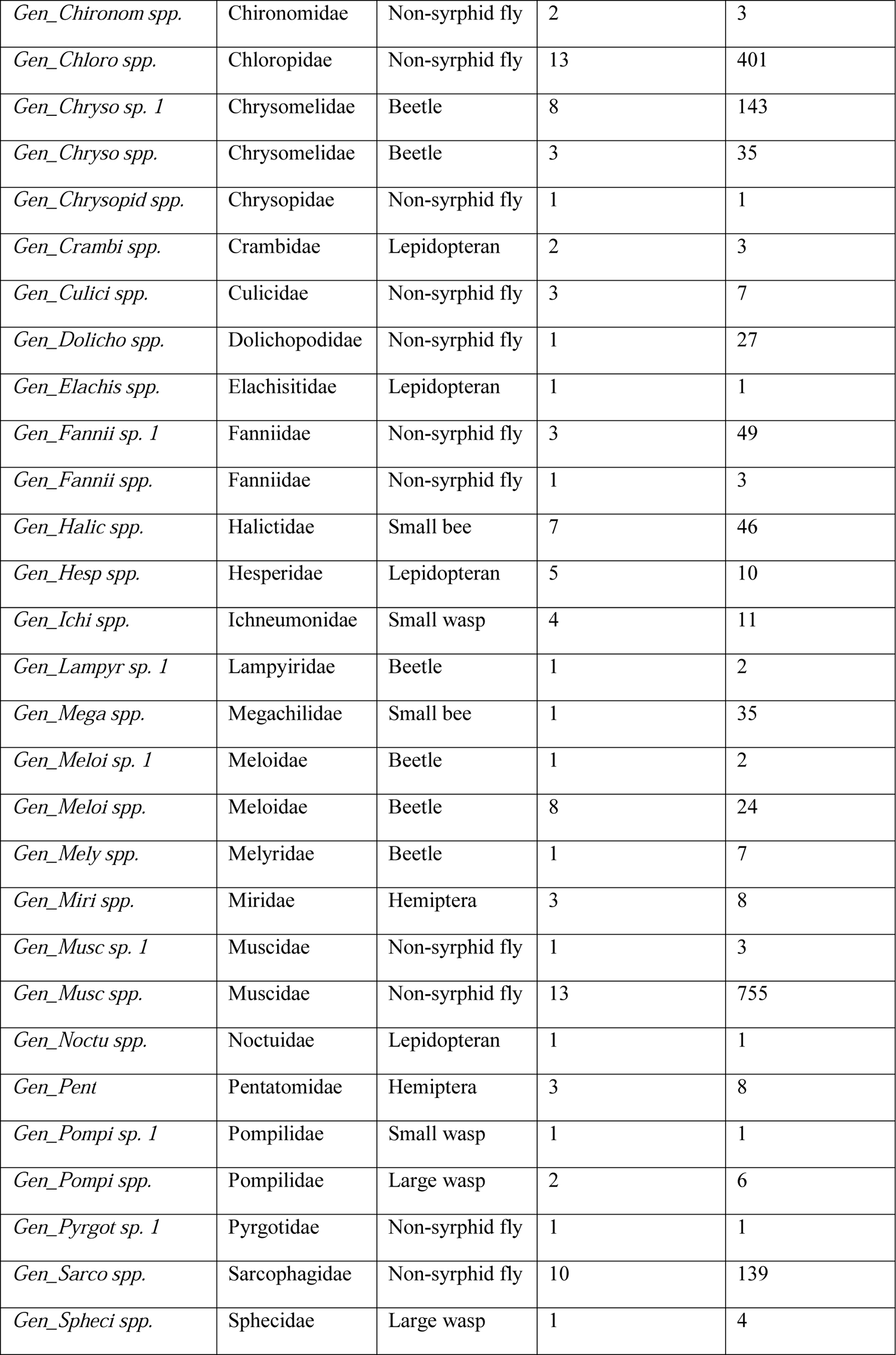
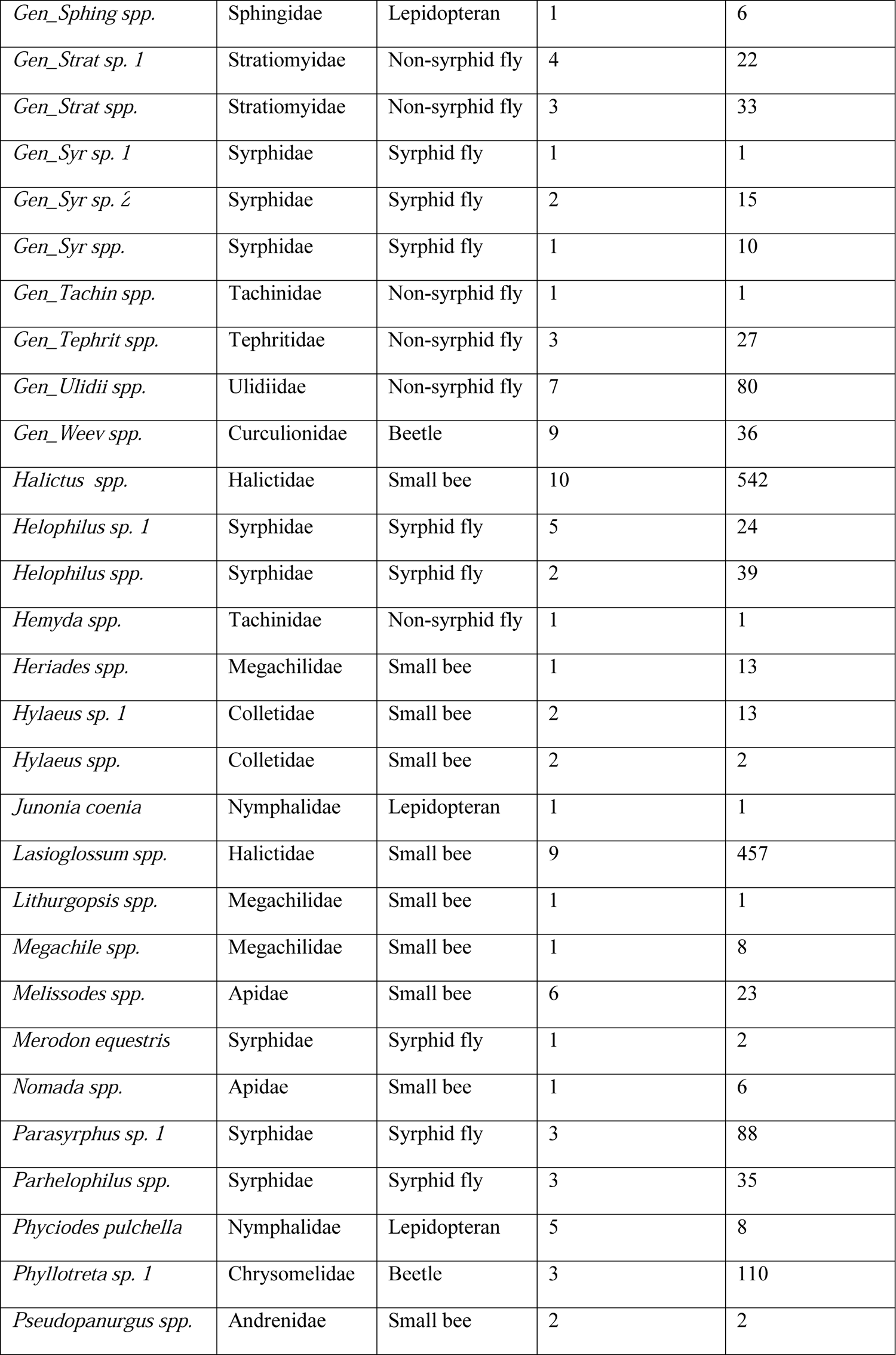
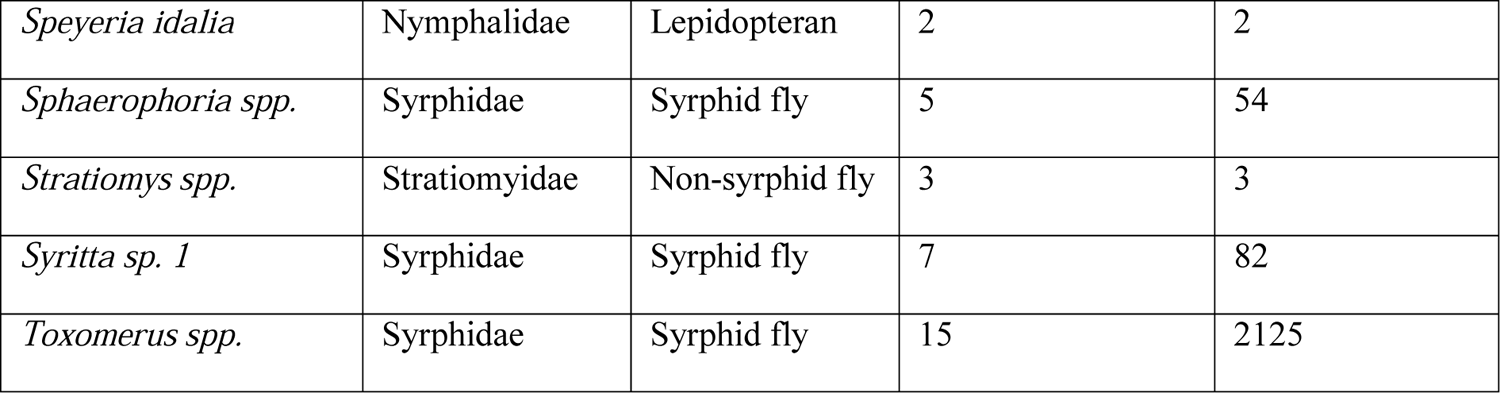

## Appendix B

The table below lists all of the biotically-pollinated flowering plants identified in the Prairie Coteau region near Brookings, South Dakota in 2019 down to lowest taxonomic level. All species except *Agrimonia sp.* was identified to species level. Plant code format is derived from United States Department of Agriculture plant database format. Average sugar contribution refers to average nectar sugar each plant species contributed per transect based on amount of sugar per inflorescence and number of inflorescences.

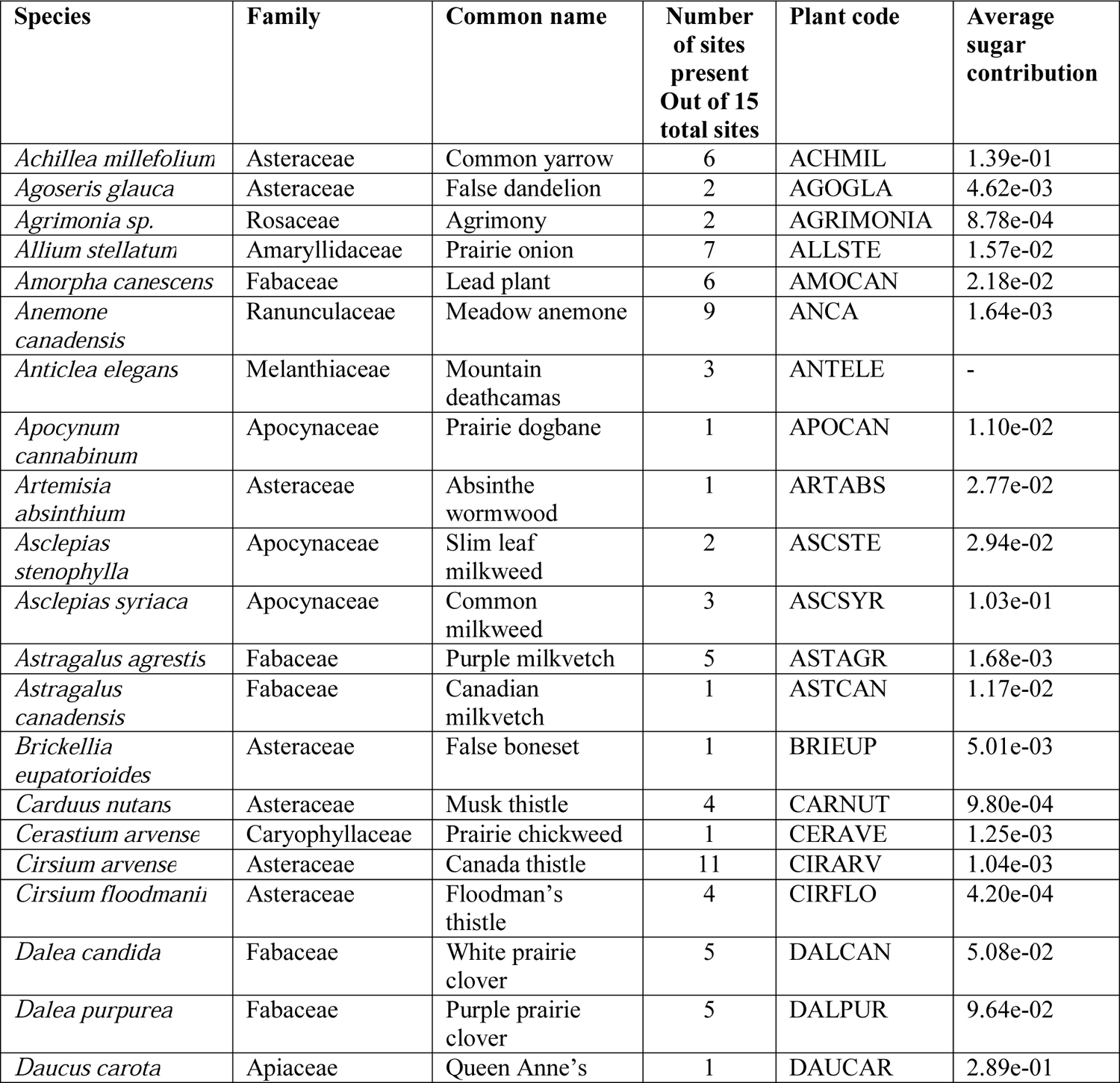
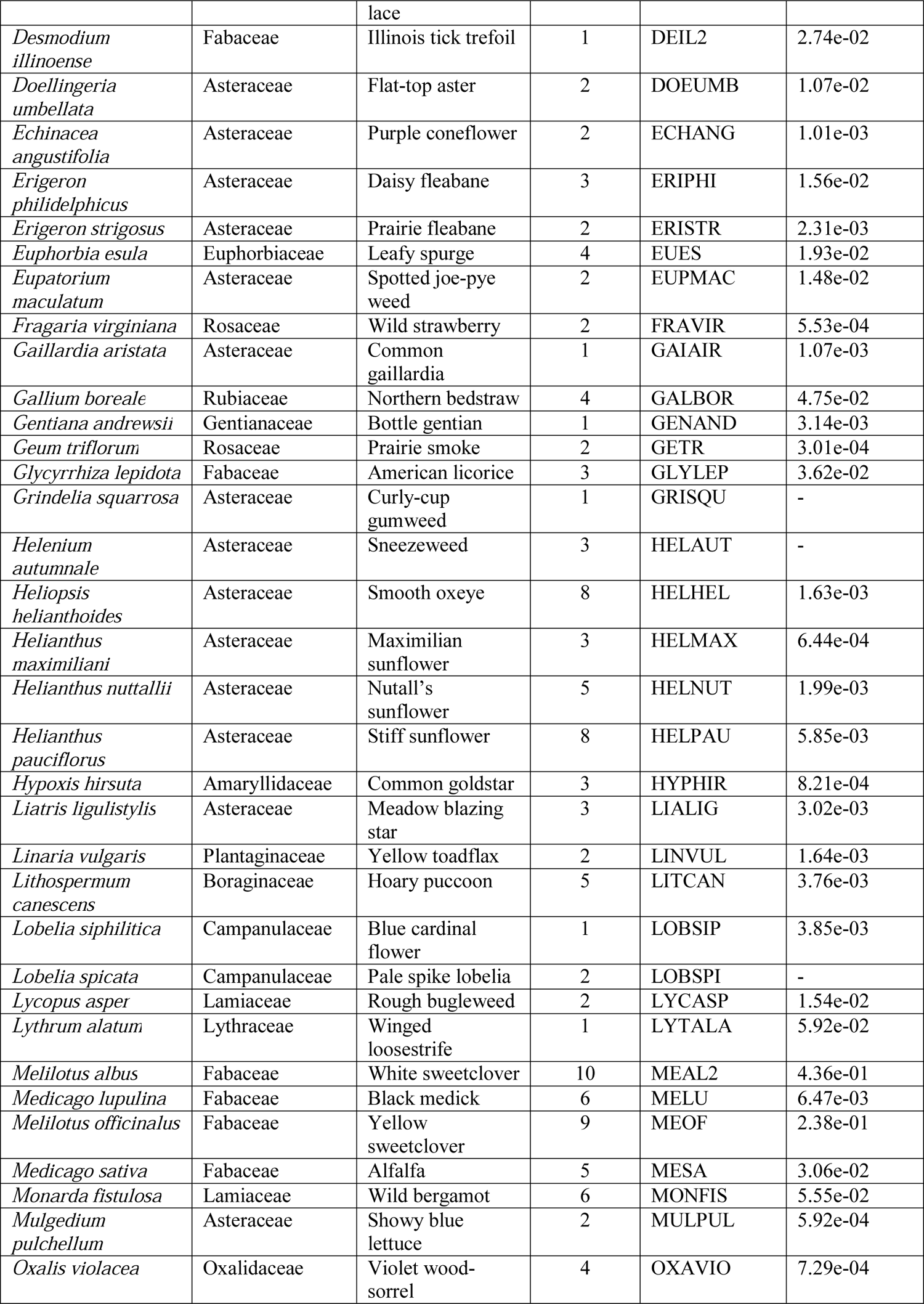
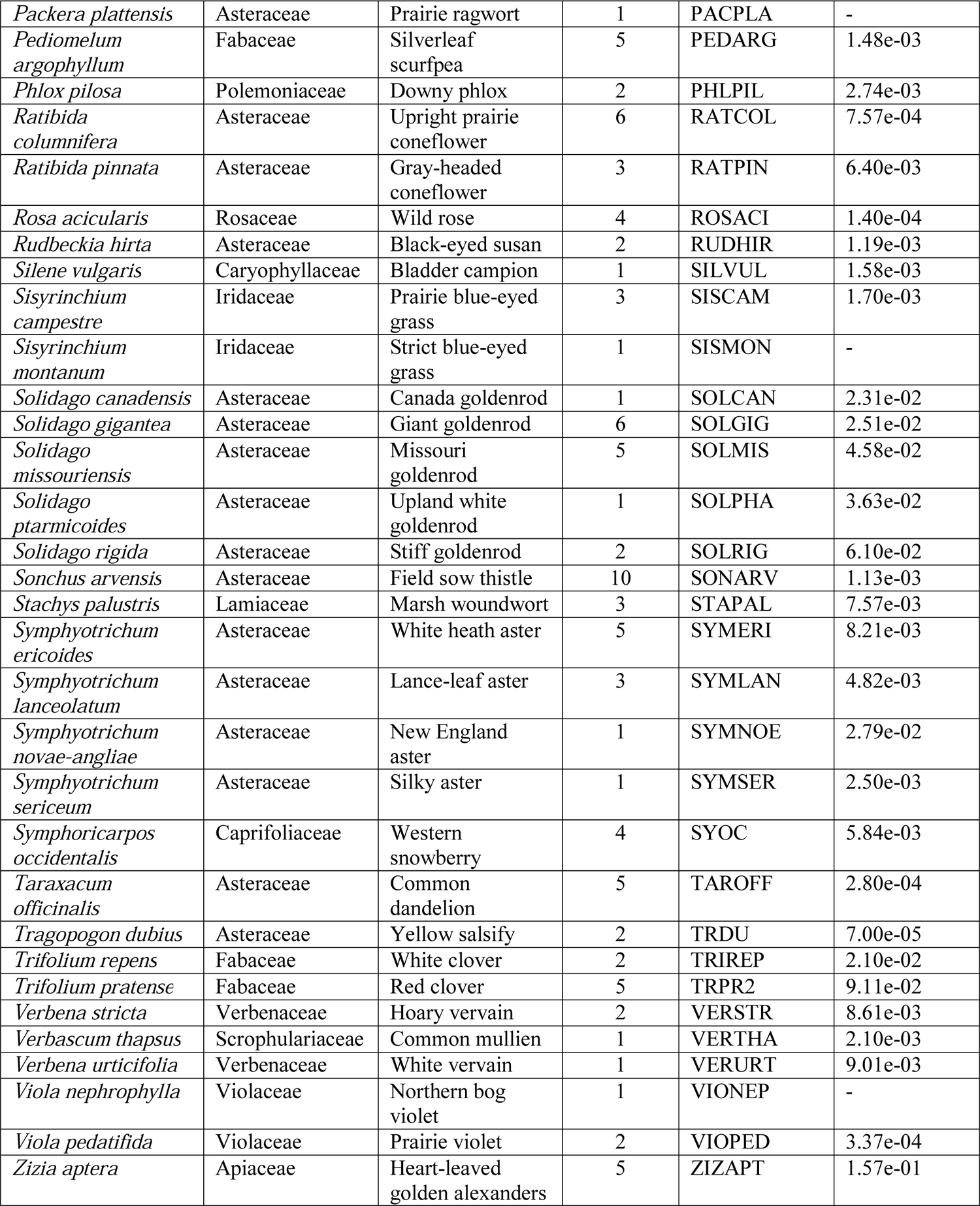

## Literature Cited

Aizen, M. A., and P. Feinsinger. 2003. Bees not to be? Responses of insect pollinator faunas and flower pollination to habitat fragmentation. In G. A. Bradshaw and P. A. Marquet [eds.], How landscapes change, 111–129. Springer-Verlag Berlin, Germany. doi:10.1007/978-3-662-05238-9_7

Aizen, M. A., C. L. Morales, and J. M. Morales. 2008. Invasive mutualists erode native pollination webs. PloS Biology, 6:396–403. doi:10.1371/journal.pbio.0060031

Alarcón, R., N. M. Waser, and J. Ollerton. 2008. Year-to-year variation in the topology of a plant – pollinator interaction network. Oikos, 117:1796–1807. doi:10.1111/j.0030-1299.2008.16987.x

Alaux, C., J. L. Brunet, C. Dussaubat, F. Mondet, S. Tchamitchan, M. Cousin, J. Brillard, A. Baldy, L. P. Belzunces, and Y. Le Conte. 2010. Interactions between Nosema microspores and a neonicotinoid weaken honey bees (*Apis mellifera*). Environmental Microbiology, 12:774–782. doi:10.1111/j.1462-2920.2009.02123.x

Albrecht, M., P. Duelli, C. Müller, D. Kleijn, and B. Schmid. 2007. The Swiss agri□environment scheme enhances pollinator diversity and plant reproductive success in nearby intensively managed farmland. Journal of Applied Ecology, 44:813–822. doi:10.1111/j.1365-2664.2007.01306.x

Albrecht, M., M. Riesen, and B. Schmid. 2010. Plant – pollinator network assembly along the chronosequence of a glacier foreland. Oikos, 119:1610–1624. Doi: 10.1111/j.1600-0706.2010.18376.x

Albrecht, M., B. Schmid, Y. Hautier, and C. B. Müller. 2012. Diverse pollinator communities enhance plant reproductive success. Proceedings of the Royal Society B: Biological Sciences, 279:4845–4852. Doi: 10.1098/rspb.2012.1621

Arceo-Gómez, G., D. Barker, A. Stanley, T. Watson, and J. Daniels. 2020. Plant-pollinator network structural properties differentially affect pollen transfer dynamics and pollination success. Oecologia, 192:1037–1045. doi:10.1007/s00442-020-04637-5

Ashworth, L., R. Aguilar, L. Galetto, and M. A. Aizen. 2004. Why do pollination generalist and specialist plant species show similar reproductive susceptibility to habitat fragmentation? Journal of Ecology, 92:717–719. Doi: 10.1111/j.0022-0477.2004.00910.x

Bartomeus, I., V. Montserrat, and L. Santamaria. 2008. Contrasting effects of invasive plants in plant-pollinator networks. Oecologia, 155:761–770. doi:10.1007/s00442-007-0946-1

Bartomeus, I., J. S. Ascher, D. Wagner, B. N. Danforth, S. Colla, S. Kornbluth, and R. Winfree. 2011. Climate-associated phenological advances in bee pollinators and bee-pollinated plants. Proceedings of the National Academy of Sciences, 108:20645–20649. doi:10.1073/pnas.1115559108

Bascompte, J., P. Jordano, C. J. Melián, and J. M. Olesen. 2003. The nested assembly of plantanimal mutualistic networks. Proceedings of the National Academy of Sciences, 100:9383–9387. doi:10.1073/pnas.1633576100

Bascompte, J., P. Jordano, and J. M. Olesen. 2006. Asymmetric coevolutionary networks facilitate biodiversity maintenance. Science, 312:431–433. Doi: 10.1126/science.1123412

Bates, D., M. Maechler, B. Bolker, and S. Walker. 2015. Fitting Linear Mixed-Effects Models Using lme4. Journal of Statistical Software, 67:1–48. doi:10.18637/jss.v067.i01.

Bauman, P., J. Blastick, C. Grewing, and A. Smart. 2016. Quantifying undisturbed land on South Dakota’s Prairie Coteau. A report to The Nature Conservancy from South Dakota State University based on the Prairie Coteau boundary as defined by the April 30, 2010 TNC National Fish and Wildlife Foundation Business Plan “Conserving and Restoring Tallgrass Prairie: Prairie Coteau, South Dakota and Minnesota”.

Beekman, M., and F. L. W. Ratnieks. 2000. Long-range foraging by the honeybee, *Apis mellifera L*. Functional Ecology, 14:490–496. doi:10.1046/j.1365-2435.2000.00443.x

Begosh, A., L. M. Smith, S. T. McMurry, and J. P. Harris. 2020. Influence of the Conservation Reserve Program (CRP) and playa wetlands on pollinator communities in the Southern High Plains, USA. Journal of Environmental Management, 256, 109910. doi:10.1016/j.jenvman.2019.109910

Benton, T. G., J. A. Vickery, and J. D. Wilson. 2003. Farmland biodiversity: is habitat heterogeneity the key? Trends in Ecology & Evolution, 18:182–188. doi:10.1016/s0169-5347(03)00011-9

Bivand, R., and C. Rundel. 2020a. rgeos: Interface to Geometry Engine -Open Source (‘GEOS’). R package version 0.5–5.https://CRAN.R-project.org/package=rgeos

Bivand, R., T. Keitt and B. Rowlingson. 2020b. Rgdal: Bindings for the ‘Geospatial’ Data Abstraction Library. R package version 1.5–18.https://CRAN.R-project.org/package=rgdal

Bivand, R., and N. Lewin-Koh. 2020c. Maptools: Tools for Handling Spatial Objects. R package version 1.0–2. https://CRAN.R-project.org/package=maptools

Blüthgen, N., F. Menzel, and N. Blüthgen. 2006. Measuring specialization in species interaction networks. BMC ecology, 6:9. doi:10.1186/1472-6785-6-9

Blüthgen, N., J. Fründ, D. P. Vázquez, and F. Menzel. 2008. What do interaction network metrics tell us about specialization and biological traits. Ecology, 89:3387–3399. doi:10.1890/07-2121.1

Bock, C. E., Z. F. Jones, and J. H. Bock. 2008. The oasis effect: response of birds to exurban development in a southwestern savanna. Ecological Applications, 18:1093–1106. doi:10.1890/07-1689.1

Boyle, N. K., and T. L. Pitts-Singer. 2019. Assessing blue orchard bee *(Osmia lignaria)* propagation and pollination services in the presence of honey bees *(Apis mellifera)* in Utah tart cherries. PeerJ, 7:e7639. doi:10.7717/peerj.7639

Brown, B.J., R. J. Mitchell, and S. A. Graham. 2002. Competition for pollination between an invasive species (purple loosestrife) and a native congener. Ecology, 83:2328–2336. doi:10.1890/0012-9658(2002)083[2328:cfpbai]2.0.co;2

Burkle, L. A., and R. Alarcón. 2011. The future of plant-pollinator diversity: understanding interaction networks across time, space, and global change. American Journal of Botany, 98:528–538. doi:10.3732/ajb.1000391

Burkle, L., and R. Irwin. 2009. The importance of interannual variation and bottom-up nitrogen enrichment for plant – pollinator networks. Oikos, 118:1816–1829. Doi: 10.1111/j.1600-0706.2009.17740.x,

Calinger, K. M., S. Queenborough, and P. S. Curtis. 2013. Herbarium specimens reveal the footprint of climate change on flowering trends across north-central North America. Ecology Letters, 16:1037–1044. Doi: 10.1111/ele.12135

Campbell, D.R. 1985. Pollinator sharing and seed set of Stellaria pubera: competition for pollination. Ecology, 66:544–553. doi:10.2307/1940403

CaraDonna, P. J., A. M. Iler, and D. W. Inouye. 2014. Shifts in flowering phenology reshape a subalpine plant community. Proceedings of the National Academy of Sciences, USA 111: 4916–4921. doi:10.1073/pnas.1323073111

Carvalheiro, L. G., C. L. Seymour, R. Veldtman, and S. W. Nicolson. 2010. Pollination services decline with distance from natural habitat even in biodiversity□rich areas. Journal of Applied Ecology, 47:810–820. doi:10.1111/j.1365-2664.2010.01829.x

Chiari, W.C., V.D.A.A.D. Toledo, M.C.C. Ruvolo-Takasusuki, A.J.B.D. Oliveira, E.S. Sakaguti, V.M. Attencia, F.M. Costa, and M.H. Mitsui. 2005. Pollination of soybean (Glycine max L. Merril) by honeybees *(Apis mellifera L.)*. Brazilian archives of biology and technology, 48:31–36. doi:10.1590/s1516-89132005000100005

Costanza, R., R. d’Arge, R. De Groot, S. Farber, M. Grasso, B. Hannon, K. Limburg, S. Naeem, R.V. O’Neill, J. Paruelo and R.G. Raskin. 1998. The value of ecosystem services: putting the issues in perspective. Ecological economics, 25:67–72.

Cox-Foster, D.L., S. Conlan, E. C. Holmes, … D. M. Geiser, and V. Martinson. 2007. A metagenomic survey of microbes in honeybee colony collapse disorder. Science, 318:283–287. doi:10.1126/science.1146498

Cox-Foster, D. L., and D. VanEngelsdorp. 2009. Saving the honeybee. Scientific American, 300:40–47. doi:10.1038/SCIENTIFICAMERICAN0409-40

Cunningham-Minnick, M.J., Peters, V.E. and Crist, T.O., 2019. Nesting habitat enhancement for wild bees within soybean fields increases crop production. Apidologie, 50:833–844. doi:10.1007/s13592-019-00691-y

Cunningham-Minnick, M. J., and T. O. Crist. 2020. Floral resources of an invasive shrub alter native bee communities at different vertical strata in forest-edge habitat. Biological Invasions, 22:2283–2298. doi:10.1007/s10530-020-02248-y

Cusser, S., and K. Goodell. 2013. Diversity and distribution of floral resources influence the restoration of plant-pollinator networks on a reclaimed strip mine. Restoration Ecology, 21:713–721. doi:10.1111/rec.12003

Dormann, C. F., J. Fruend, B. Gruber, and M. C. F. Dormann. 2009. Package bipartite. version 2.15

Dunne, J. A., R. J. Williams, and N. D. Martinez. 2002. Food-web structure and network theory: the role of connectance and size. Proceedings of the National Academy of Sciences, 99:12917–12922. doi:10.1073/pnas.192407699

Dupont, Y. L., and J. M. Olesen. 2009a. Ecological modules and roles of species in heathland plant-insect flower visitor networks. Journal of Animal Ecology, 78:346–353. doi:10.11/j.l365-2656.2008.01501.x

Dupont, Y. L., B. Padrón, J. M. Olesen, and T. Petanidou. 2009b. Spatio-temporal variation in the structure of pollination networks. Oikos, 118:1261–1269. Doi: 10.1111/j.1600-0706.2009.17594.x

Fahrig, L., J. Baudry, L. Brotons, F. G. Burel, T. O. Crist, R. J. Fuller, C. Sirami, G. M. Siriwardena, and J. L Martin. 2011. Functional landscape heterogeneity and animal biodiversity in agricultural landscapes. Ecology Letters, 14:101–112. doi:10.1111/j.1461-0248.2010.01559.x

Fenster, C. B., W. S. Armbruster, P. Wilson, M. R. Dudash, and J. D. Thomson. 2004. Pollination syndromes and floral specialization. Annu. Rev. Ecol. Evol. Syst., 35:375–403. doi:10.1146/annurev.ecolsys.34.011802.132347

Ferreira, P. A., D. Boscolo, and B. F. Viana. 2013. What do we know about the effects of landscape changes on plant-pollinator interaction networks? Ecological Indicators, 31:35–40. doi:10.1016/j.ecolind.2012.07.025

Fitter, A. H. and R. S. R. Fitter. 2002. Rapid Changes in Flowering Time in British Plants. Science, 296:1689–1691. doi:10.1126/science.1071617

Foley, J. A., R. DeFries, G. P. Asner, … N. Ramankutty, and P. K. Snyder. 2005. Global consequences of land use. Science, 309:570–574. doi:10.1126/science.1111772

Fontaine, C., I. Dajoz, J. Meriguet, and M. Loreau. 2005. Functional diversity of plant-pollinator interaction webs enhances the persistence of plant communities. PLoS Biology, 4:e1. doi:10.1371/journal.pbio.0040001

Forister, M. L., and A. M. Shapiro. 2003. Climatic trends and advancing spring flight of butterflies in lowland California. Global Change Biology 9: 1130–1135. doi:10.1046/j.1365-2486.2003.00643.x

Fowler, R.E., E. L. Rotheray, and D. Goulson. 2016. Floral abundance and resource quality influence pollinator choice. Insect Conservation and Diversity, 9:481–494.

Fründ, J., K. E. Linsenmair, and N. Blüthgen. 2010. Pollinator diversity and specialization in relation to flower diversity. Oikos, 119:1581–1590. doi:10.1111/j.1600-0706.2010.18450.x

Garibaldi, L. A., I. Steffan□Dewenter, C. Kremen, … S. S. Greenleaf, and A. Holzschuh. 2011. Stability of pollination services decreases with isolation from natural areas despite honeybee visits. Ecology Letters, 14:1062–1072. doi:10.1111/j.1461-0248.2011.01669.x

Gotlieb, A., Y. Hollender, and Y. Mandelik. 2011. Gardening in the desert changes bee communities and pollination network characteristics. Basic and Applied Ecology, 12:310–320. doi:10.1016/j.baae.2010.12.003

Greer, M. J., K. K. Bakker, and C. D. Dieter. 2016. Grassland bird response to recent loss and degradation of native prairie in central and western South Dakota. The Wilson Journal of Ornithology, 128:278–289. doi:10.1676/wils-128-02-278-289.1

Habel, J. C., W. Ulrich, N. Biburger, S. Seibold, and T. Schmitt. 2019. Agricultural intensification drives butterfly decline. The Royal Entomological Society, 12:289–295. doi:10.1111/icad.12343

Hadley, A. S., and M. G. Betts. 2012. The effects of landscape fragmentation on pollination dynamics: absence of evidence not evidence of absence. Biological Reviews, 87:526–544. doi:10.1111/j.1469-185x.2011.00205.x

Han, W., Z. Yang, L. Di, and R. Mueller. 2012. CropScape: A Web service based application for exploring and disseminating US conterminous geospatial cropland data products for decision support. Computers and Electronics in Agriculture, 84, 111–123.

Hass, A. L., L. Brachmann, P. Batáry, Y. Clough, H. Behling, and T. Tscharntke. 2019. Maize□dominated landscapes reduce bumblebee colony growth through pollen diversity loss. Journal of Applied Ecology, 56:294–304. doi:10.1111/1365-2664.13296

Heard, M. S., C. Carvell, N. L. Carreck, P. Rothery, J. L. Osborne, and A. F. G. Bourke. 2007. Landscape context not patch size determines bumble-bee density on flower mixtures sown for agri-environment schemes. Biology Letters, 3:638–641. doi:10.1098/rsbl.2007.0425

Hernández□Castellano, C., A. Rodrigo, J. M. Gómez, C. Stefanescu, J. A. Calleja, S. Reverté, and J. Bosch. 2020. A new native plant in the neighborhood: effects on plant-pollinator networks, pollination, and plant reproductive success. Ecology, 101:e03046. doi:10.1002/ecy.3046

Hinners, S. J., and M. K. Hjelmroos-Koski. 2009. Receptiveness of foraging wild bees to exotic landscape elements. The American Midland Naturalist, 162:253–265. doi:10.1674/0003-0031-162.2.253

Hoehn, P., T. Tscharntke, J. M. Tylianakis, and I. Steffan-Dewenter. 2008. Functional group diversity of bee pollinators increases crop yield. Proceedings of the Royal Society B: Biological Sciences, 275:2283–2291. doi:10.1098/rspb.2008.0405

Hoekstra, J. M., T. M. Boucher, T. H. Ricketts, and C. Roberts. 2005. Confronting a biome crisis: global disparities of habitat loss and protection. Ecology letters, 8:23–29. doi:10.1111/j.1461-0248.2004.00686.x

Jauker, F., B. Jauker, I. Grass, I. Steffan□Dewenter, and V. Wolters. 2019. Partitioning wild bee and hoverfly contributions to plant-pollinator network structure in fragmented habitats. Ecology, 100:e02569. doi:10.1002/ecy.2569

Jordano, P., J. Bascompte, and J. M. Olesen. 2003. Invariant properties in coevolutionary networks of plant-animal interactions. Ecology Letters, 6:69–81. doi:10.1046/j.1461-0248.2003.00403.x

Kandori, I., T. Hirao, S. Matsunaga, and T. Kurosaki. 2009. An invasive dandelion unilaterally reduces the reproduction of a native congener through competition for pollination. Oecologia, 159:559–569.

Kearns, C. A., D. W. Inouye, and N. M. Waser. 1998. Endangered mutualisms: the conservation of plant-pollinator interactions. Annu. Rev. Ecol. Evol. Syst, 29:83–112. doi:10.1146/annurev.ecolsys.29.1.83

Klecka, J., J. Hadrava, P. Biella, and A. Akter. 2018. Flower visitation by hoverflies (Diptera: Syrphidae) in a temperate plant-pollinator network. PeerJ, 6:e6025. doi:10.7717/peerj.6025

Klein, A., B. E. Vaissiere, J. H. Cane, I. Steffan-Dewenter, S. A. Cunningham, C. Kremen, and T. Tscharntke. 2007. Importance of pollinators in changing landscapes for world crops. Proceedings of the Royal Society, 274:303–313. doi:10.1098/rspb.2006.3721

Knight, M. E., J. L. Osborne, R. A. Sanderson, R. J. Hale, A. P. Martin, and D. Goulson. 2009. Bumblebee nest density and the scale of available forage in arable landscapes. Insect Conservation and Diversity, 2:116–124. doi:10.1111/j.1752-4598.2009.00049.x

Kremen, C., N. M. Williams, and R. W. Thorp. 2002. Crop pollination from native bees at risk from agricultural intensification. Proceedings of the National Academy of Sciences, 99:16812–16816. doi:10.1073/pnas.262413599

Kremen, C., N. M. Williams, R. L. Bugg, J. P. Fay, and R. W. Thorp. 2004. The area requirements of an ecosystem service: crop pollination by native bee communities in California. Ecology Letters, 7:1109–1119. doi:10.1111/j.1461-0248.2004.00662.x

Larson, D.L., P.A. Rabie, S. Droege, J. L Larson, and M. Haar. 2016. Exotic plant infestation is associated with decreased modularity and increased numbers of connectors in mixed-grass prairie pollination networks. PloS One, 11:e0155068. doi:10.1371/journal.pone.0155068

Lautenbach, S., R. Seppelt, J. Liebscher, and C.F. Dormann. 2012. Spatial and temporal trends of global pollination benefit. PLoS One, 7:e35954. doi:10.1371/journal.pone.0035954

Lázaro, A., F. Fuster, D. Alomar, and Ø. Totland. 2020. Disentangling direct and indirect effects of habitat fragmentation on wild plants’ pollinator visits and seed production. Ecological Applications, e02099. doi:10.1002/eap.2099

Liu, Z., Y. Liu, and M. H. A. Baig. 2019. Biophysical effect of conversion from croplands to grasslands in water-limited temperate regions of China. The Science of the Total Environment, 648:315–324. doi:10.1016/j.scitotenv.2018.08.128

Lopezaraiza-Mikel, M. E., R. B. Hayes, M. R. Whalley, and J. Memmott. 2007. The impact of an alien plant on a native plant-pollinator network: an experimental approach. Ecology, Letters 10:539–550. doi:10.1111/j.1461-0248.2007.01055.x

Lundgren, J. G., and S. W. Fausti. 2015. Trading biodiversity for pest problems. Science Advances, 1: e1500558. doi:10.1126/sciadv.1500558

Mallinger, R.E., J. Bradshaw, A.J. Varenhorst, and J.R. Prasifka. 2019. Native solitary bees provide economically significant pollination services to confection sunflowers (*Helianthus annuus L*.)(Asterales: Asteraceae) grown across the Northern Great Plains. Journal of Economic Entomology, 112:40–48. doi:10.1093/jee/toy322

Memmott, J., N. M. Waser, and M. V. Price. 2004. Tolerance of pollination networks to species extinctions. Proceedings of the Royal Society of London. Series B: Biological Sciences, 271:2605–2611. doi:10.1098/rspb.2004.2909

Milfont, M.D.O., E.E.M Rocha, A.O.N. Lima, and B.M. Freitas. 2013. Higher soybean production using honeybee and wild pollinators, a sustainable alternative to pesticides and autopollination. Environmental Chemistry Letters, 11:335–341. doi:10.1007/s10311-013-0412-8

Miranda, G. F. G., A. D. Young, M. M. Locke, S. A. Marshall, J. H. Skevington, and F. C. Thompson. 2013. Key to the genera of Nearctic Syrphidae. Canadian Journal of Arthropod Identification, 23:351. doi:10.3752/cjai.2013.23

Montoya, J. M., and G. Yvon-Durocher. 2007. Ecological networks: information theory meets Darwin’s entangled bank. Current Biology, 17:128–130. doi:10.1016/j.cub.2007.01.028

Morales, C.L. and A. Traveset. 2008. Interspecific pollen transfer: magnitude, prevalence and consequences for plant fitness. Critical Reviews in Plant Sciences, 27:221–238. doi:10.1080/07352680802205631

Moreira, E. F., D. Boscolo, and B. F. Viana. 2015. Spatial heterogeneity regulates plant-pollinator networks across multiple landscape scales. PloS One, 10:e0123628. doi:10.1371/journal.pone.0123628

Morris, W. F. 1993. Predicting the Consequence of Plant Spacing and Biased Movement for 754 Pollen Dispersal by Honey bees. Ecology, 74:493–500. doi:10.2307/1939310

Morton, E.M. and N.E. Rafferty. 2017. Plant-pollinator interactions under climate change: The use of spatial and temporal transplants. Applications in Plant Sciences, 5:1600133.

Naug, D. 2009. Nutritional stress due to habitat loss may explain recent honeybee colony collapses. Biological Conservation, 142:2369–2372. doi:10.1016/j.biocon.2009.04.007

Nielsen, A., and J. Bascompte. 2007. Ecological networks, nestedness and sampling effort. Journal of Ecology, 95:1134–1141. Doi: 10.11111/j.1365-2745.2007.01271.

Nottebrock, H., B. Schmid, K. Mayer, C. Devaux, K. J. Esler, K. Böhning□Gaese, M. Schleuning, J. Pagel, and F. M. Schurr. 2017. Sugar landscapes and pollinator□mediated interactions in plant communities. Ecography, 40:1129–1138. doi:10.1111/ecog.02441

Oksanen, J., F. G. Blanchet, M. Friendly, R. Kindt, P. Legendre, D. McGlinn, P. R. Minchin, R. B. O’Hara, G. L. Simpson, and H. Wagner. 2019. vegan: Community Ecology Package. R package version 2.5-6. https://CRAN.R-project.org/package=vegan

Okuyama, T., and J. N. Holland. 2008. Network structural properties mediate the stability of mutualistic communities. Ecology Letters, 11:208–216. doi:10.1111/j.1461-0248.2007.01137.x

Olesen, J. M. and S. K. Jain. 1994. D: Fragmented plant populations and their lost interactions. In V. Loeschcke, S. K. Jain, and J. Tomiuk [eds.], Conservation Genetics, 417–426. Birkhäuser, Basel. doi:10.1007/978-3-0348-8510-2

Olesen, J. M., J. Bascompte, Y. L. Dupont, and P. Jordano. 2007. The modularity of pollination networks. Proceedings of the National Academy of Sciences, 104:19891–19896. doi:10.1073/pnas.0706375104

Otto, C. R., C. L. Roth, B. L. Carlson, and M. D. Smart. 2016. Land-use change reduces habitat suitability for supporting managed honeybee colonies in the Northern Great Plains. Proceedings of the National Academy of Sciences, 113:10430–10435. doi:10.1073/pnas.1603481113

Otto, C.R.V., A. Smart, R. S. Cornman, M. Simanonok, and D.D. Iwanowicz. 2020. Forage and habitat for pollinators in the northern Great Plains—Implications for U.S. Department of Agriculture conservation programs: U.S. Geological Survey Open-File Report 20201037, 64 p. Doi: 10.3133/ofr20201037

Ouvrard, P., J. Transon, and A. L. Jacquemart. 2018. Flower-strip agri-environment schemes provide diverse and valuable summer flower resources for pollinating insects. Biodiversity and Conservation, 27:2193–2216. Doi: 10.1007/s10531-018-1531-0

Padrón, B., A. Traveset, T. Biedenweg, D. Diaz, M. Nogales, and J. M. Olesen. 2009. Impact of alien plant invaders on pollination networks in two archipelagos. PLoS One, 4:e6275. doi:10.1371/journal.pone.0006275

Perpinan, O., and R. Hijmans. 2020. rasterVis. R package version 0.49.

Petanidou, T., A. S. Kallimanis, J. Tzanopoulos, S. P. Sgardelis, and J.P. Pantis. 2008. Long-term observation of a pollination network: Fluctuation in species and interactions, relative invariance of network structure and implications for estimates of speciation. Ecology Letters, 11:564–575. Doi: 10.1111/j.1461-0248.2008.01170.x

Peterson, M.J., D.M. Hall, A.M. Feldpausch-Parker and T.R. Peterson. 2010. Obscuring ecosystem function with application of the ecosystem services concept. Conservation Biology, 24:113–119. Doi: 10.1111/j.1523-1739.2009.01305.x

Pettis, J. S., E. M. Lichtenberg, M. Andree, J. Stitzinger, and R. Rose. 2013. Crop pollination exposes honey bees to pesticides which alters their susceptibility to the gut pathogen *Nosema ceranae*. PloS One, 8:e70182. doi:10.1371/journal.pone.0070182

Pinheiro J., D. Bates, S. DebRoy, D. Sarkar, and R Core Team. 2020. nlme: Linear and Nonlinear Mixed Effects Models. R package version 3.1-144, <URL: https://CRAN.R-project.org/package=nlme>.

Ponisio, L. C., P. de Valpine, L. K. M’Gonigle, and C. Kremen. 2019. Proximity of restored hedgerows interacts with local floral diversity and species’ traits to shape long□term pollinator metacommunity dynamics. Ecology Letters, 22:1048–1060. doi:10.1111/ele.13257

Potts, S. G., B. Vulliamy, A. Dafni, G. Ne’eman, and P. Willmer. 2003. Linking bees and flowers: how do floral communities structure pollinator communities? Ecology, 84:2628–2642. doi:10.1890/02-0136

Potts, S. G., J. C. Biesmeijer, C. Kremen, P. Neumann, O. Schweiger, and W. E. Kunin. 2010. Global pollinator declines: trends, impacts and drivers. Trends in Ecology & Evolution, 25:345–353. doi:10.1016/j.tree.2010.01.007

Potts, S. G., B. Vulliamy, S. Roberts, C. O’Toole, A. Dafni, G. Ne’eman, and P. Willmer. 2005. Role of nesting resources in organizing diverse bee communities in a Mediterranean landscape. Ecological Entomology, 30:78–85. doi:10.1111/j.0307-6946.2005.00662.x

Potts, S. G., B. A. Woodcock, S. P. M. Roberts, T. Tscheulin, E. S. Pilgrim, V. K. Brown, and J. R. Tallowin. 2009. Enhancing pollinator biodiversity in intensive grasslands. Journal of Applied Ecology, 46:369–379. doi:10.1111/j.1365-2664.2009.01609.x

R Core Team. 2013. R: A language and environment for statistical computing. version 3.6.3

Ramankutty, N., and J. A. Foley. 1999. Estimating historical changes in global land cover: Croplands from 1700 to 1992. Global Biogeochemical Cycles, 13:997–1027. doi:10.1029/1999gb900046

Rashford, B. S., J. A. Walker, and C. T. Bastian. 2011. Economics of grassland conversion to cropland in the Prairie Pothole Region. Conservation Biology, 25:276–284. doi:10.1111/j.1523-1739.2010.01618

Redhead, J. W., B. A. Woodcock, M. J. Pocock, R. F. Pywell, A. J. Vanbergen, and T.H. Oliver. 2018. Potential landscape□scale pollinator networks across Great Britain: structure, stability, and influence of agricultural land cover. Ecology Letters, 21:1821–1832. doi:10.1111/ele.13157

Requier, F., K. K. Jowanowitsch, K. Kallnik, and I. Steffan□Dewenter. 2020. Limitation of complementary resources affects colony growth, foraging behavior, and reproduction in bumble bees. Ecology, 101:e02946. doi:10.1002/ecy.2946

Robinson, S. V. J., G. Losapio, and G. H. R. Henry. 2018. Flower-power: Flower diversity is a stronger predictor of network structure than insect diversity in an Arctic plant-pollinator network. Ecological complexity, 36:1–6. doi:10.1016/j.ecocom.2018.04.005

Rodríguez□Gironés, M. A., and L. Santamaria. 2006. A new algorithm to calculate the nestedness temperature of presence-absence matrices. Journal of Biogeography, 33:924–935. doi:10.1111/j.1365-2699.2006.01444

Rogers, S. R., D. R. Tarpy, and H. J. Burrack. 2014. Bee species diversity enhances productivity and stability in a perennial crop. PloS One, 9:e97307. doi:10.1371/journal.pone.0097307

Roy, D. B., and T. H. Sparks. 2000. Phenology of British butterflies and climate change. Global Change Biology, 6:407–416. doi:10.1046/j.1365-2486.2000.00322.x

Royer, F., and R. Dickinson. 1999. Weeds of the Northern US and Canada. The University of Alberta Press, Edmonton, Canada.

Sakai, S., S. Metelmann, Y. Toquenaga, and A. Telschow. 2016. Geographical variation in the heterogeneity of mutualistic networks. Royal Society Open Science, 3:150630. doi:10.1098/rsos.150630

Smart M. D., J. S. Pettis, N. H. Euliss, and M. S. Spivak. 2016. Land use in the Northern Great Plains region of the U.S. influences the survival and productivity of honeybee colonies. Agriculture, Ecosystems, and Environment, 230:139–149. doi:10.1016/j.agee.2016.05.030

Soares, R. G. S., P. A. Ferreira, and L. E. Lopes. 2017. Can plant-pollinator network metrics indicate environmental quality? Ecological Indicators, 78:361–370. doi:10.1016/j.ecolind.2017.03.037

Sommaggio, D. 1999. Syrphidae: can they be used as environmental bioindicators? Agriculture, Ecosystems & Environment, 74:343–356. doi:10.1016/s0167-8809(99)00042-0

Spellman, K.V., L. C. Schneller, C. P. Mulder, C.P. and M. L. Carlson. 2015. Effects of non-native Melilotus albus on pollination and reproduction in two boreal shrubs. Oecologia, 179:495–507. doi:10.1007/s00442-015-3364-9

Spiesman, B. J., and B. D. Inouye. 2013. Habitat loss alters the architecture of plant-pollinator interaction networks. Ecology, 94:2688–2696. doi:10.1890/13-0977.1

Spleen, A. M., E. J. Lengerich, K. Rennich, … J. Stitzinger, and K. Lee. 2013. A national survey of managed honeybee 2011-12 winter colony losses in the United States: results from the Bee Informed Partnership. Journal of Apicultural Research, 52:44–53. doi:10.3896/IBRA.1.52.2.07

Stang, M., P. G. L. Klinkhamer, and E. van der Meijden. 2006. Size constraints and flower abundance determine the number of interactions in a plant-flower visitor web. Oikos, 112:111–121. doi:10.1111/j.0030-1299.2006.14199

Steffan-Dewenter, I., U. Münzenberg, C. Bürger, C. Thies, and T. Tscharntke. 2002. Scale□dependent effects of landscape context on three pollinator guilds. □Ecology, 83:1421–1432. doi:10.1890/0012-9658(2002)083[1421:sdeolc]2.0.co;2

Steffan-Dewenter, I., S. G. Potts, and L. Packer. 2005. Pollinator diversity and crop pollination services are at risk. Trends in Ecology & Evolution, 20:651–652. doi:10.1016/j.tree.2005.09.00

Sugden, E. A., R. W. Thorp, and S. L. Buchmann. 1996. Honeybee-native bee competition: focal point for environmental change and apicultural response in Australia. Bee World, 77:26–44. doi:10.1080/0005772x.1996.11099280

Tamburini G., F. Lami, L. Marini. 2017. Pollination benefits are maximized at intermediate nutrient levels. Proc. R. Soc. B 284:20170729. doi:10.1098/rspb.2017.0729

Tepedino, V.J., B.A. Bradley, and T.L. Griswold. 2008. Might flowers of invasive plants increase native bee carrying capacity? Intimations from Capitol Reef National Park, Utah. Natural Areas Journal, 28:44–50. Doi: 10.3375/0885-8608(2008)28[44:MFOIPI]2.0.CO;2

Thébault, E., and C. Fontaine. 2010. Stability of ecological communities and the architecture of mutualistic and trophic networks. Science, 329:853–856. Doi: 10.1126/science.1188321

Thomson, D. 2004. Competitive interactions between the invasive European honeybee and native bumble bees. Ecology, 85:458–470. doi:10.1890/02-0626

Tscharntke, T., A. M. Klein, A. Kruess, I. Steffan□Dewenter, and C. Thies. 2005. Landscape perspectives on agricultural intensification and biodiversity-ecosystem service management. Ecology Letters, 8:857–874. Doi: 10.1111/j.1461-0248.2005.00782

Twerd, L., and W. Banaszak-Cibicka. 2019. Wastelands: their attractiveness and importance for preserving the diversity of wild bees in urban areas. Journal of Insect Conservation, 23:573–588. doi:10.1007/s10841-019-00148-8

Tylianakis, J. M., and R. J. Morris. 2017. Ecological networks across environmental gradients. Annual Review of Ecology, Evolution, and Systematics, 48:25–48. doi:10.1146/annurev-ecolsys-110316-022821

US Department of Agriculture, National Agricultural Statistics Service. 2014. US Department of Agriculture, Washington, DC. Available at http://usda.mannlib.cornell.edu/MannUsda/viewDocumentInfo.do?documentID=1191.

USDA National Agricultural Statistics Service Cropland Data Layer. 2019. Published crop-specific data layer [Online]. Available at https://nassgeodata.gmu.edu/CropScape/ (accessed in 2019; verified in 2019. USDA-NASS, Washington, DC.

Valdovinos, F. S., B. J. Brosi, H. M. Briggs, P. Moisset de Espanés, R. Ramos-Jiliberto, and N. D. Martinez. 2016. Niche partitioning due to adaptive foraging reverses effects of nestedness and connectance on pollination network stability. Ecology Letters, 19:1277–1286. doi:10.1111/ele.12664

Van Bruggen, T. 1985. The Vascular Plants of South Dakota. Second edition. Iowa State University Press, Ames, Iowa, USA.

Vanbergen, A. J., and the Insect Pollinators Initiative. 2013. Threats to an ecosystem service: pressures on pollinators. Frontiers in Ecology and the Environment, 11:251–259. doi:10.1890/120126

Vanbergen, A.J. 2014. Landscape alteration and habitat modification: impacts on plant-pollinator systems. Current Opinion in Insect Science, 5:44–49.

Venables, W. N. and B. D. Ripley. 2002. Modern Applied Statistics with S. Fourth Edition. Springer, New York. ISBN 0-387-95457-0

Vickruck, J. L., L. R. Best, M. P. Gavin, J. H. Devries, and P. Galpern. 2019. Pothole wetlands provide reservoir habitat for native bees in prairie croplands. Biological Conservation, 232:43–50. doi:10.1016/j.biocon.2019.01.015

Vinson S.B., G. W. Frankie, and J. Barthell. 1993. Threats to the diversity of solitary bees in a neotropical dry forest in Central America. See Ref. 121a:53–82

Waser, N.M. 1983. Competition for pollination and floral character differences among sympatric plant species: a review of evidence. In C. E. Jones and R. J. Little [eds.], Handbook of experimental pollination biology, 277–293.

Weiner, C.N., M. Werner, K. E. Linsenmair, and N. Blüthgen. 2014. Land□use impacts on plant-pollinator networks: interaction strength and specialization predict pollinator declines. Ecology, 95:466–474. doi:10.1890/13-0436.1

Williams, I. H. 2002. Insect pollination and crop production: a European perspective. □In Pollinating Bees-The Conservation Link Between Agriculture and Nature, 59–65. Ministry of Environment, Brasilia, Brazil.

Williams, N. M., D. Cariveau, R. Winfree, and C. Kremen. 2011. Bees in disturbed habitats use, but do not prefer, alien plants. Basic and Applied Ecology, 12:332–341. doi:10.1016/j.baae.2010.11.008

Winfree, R., T. Griswold, and C. Kremen. 2007. Effect of human disturbance on bee communities in a forested ecosystem. Conservation Biology, 21:213–223. doi:10.1111/j.1523-1739.2006.00574

Wright, C. K., and M. C. Wimberly. 2013. Recent land use change in the Western Corn Belt threatens grasslands and wetlands. Proceedings of the National Academy of Sciences, 110:4134–4139. doi:10.1073/pnas.1215404110

