## Supplementary Materials Figures and Tables for "Quantifying habitat and landscape effects on composition and structure of plant-pollinator networks in the US Northern Great Plains" (from another version of this preprint: version 2, 2021-02-14)

### Supplementary figure legends for: Quantifying habitat and landscape effects on composition and structure of plant-pollinator networks in the US Northern Great Plains

Fig. 1 Shannon diversity of pollinators at the functional, family and genus level in the early season (May - June 2019) in the Prairie Coteau region near Brookings, South Dakota. Site abbreviations are as follows: DC = Deer Creek, AG = Altamont GPA, AR = Aurora, BP = Brookings Prairie, OL = Oak Lake Field Station, S7 = Seven-mile Fen, SV = Severson WMA, GG = Gary Gulch, CP = Coteau Prairie WMA, CL = Coteau Lakes GPA, RBH2 = Round Bullhead Lakes Site 2, SP = Sioux Prairie, AP = Altamont Prairie, RBH1 = Round Bullhead Site 1, J = Jacobsen Fen.

Fig. 2 Shannon diversity of pollinators at the functional, family and genus level in the mid-season (July-August 2019) in the Prairie Coteau region near Brookings, South Dakota. Site abbreviations are as follows: DC = Deer Creek, AG = Altamont GPA, AR = Aurora, BP = Brookings Prairie, OL = Oak Lake Field Station, S7 = Seven-mile Fen, SV = Severson WMA, GG = Gary Gulch, CP = Coteau Prairie WMA, CL = Coteau Lakes GPA, RBH2 = Round Bullhead Lakes Site 2, SP = Sioux Prairie, AP = Altamont Prairie, RBH1 = Round Bullhead Site 1, J = Jacobsen Fen.

Fig. 3 Shannon diversity of pollinators at the functional, family and genus level in the late season (September – October 2019) in the Prairie Coteau region near Brookings, South Dakota. Prairies were only sampled once this season, thus are represented by a single bar in this graph. Site abbreviations are as follows: DC = Deer Creek, AG = Altamont GPA, AR = Aurora, BP = Brookings Prairie, OL = Oak Lake Field Station, S7 = Seven-mile Fen, SV = Severson WMA, GG = Gary Gulch, CP = Coteau Prairie WMA, CL = Coteau Lakes GPA, RBH2 = Round Bullhead Lakes Site 2, SP = Sioux Prairie, AP = Altamont Prairie, RBH1 = Round Bullhead Site 1, J = Jacobsen Fen.

Fig. 4 Shannon diversity of biotically-pollinated plants at the family, genus, and species level in the early season (May-June 2019) in the Prairie Coteau region near Brookings, South Dakota. Site abbreviations are as follows: DC = Deer Creek, AG = Altamont GPA, AR = Aurora, BP = Brookings Prairie, OL = Oak Lake Field Station, S7 = Seven-mile Fen, SV = Severson WMA, GG = Gary Gulch, CP = Coteau Prairie WMA, CL = Coteau Lakes GPA, RBH2 = Round Bullhead Lakes Site 2, SP = Sioux Prairie, AP = Altamont Prairie, RBH1 = Round Bullhead Site 1, J = Jacobsen Fen.

Fig. 5 Shannon diversity of biotically-pollinated plants at the family, genus, and species level during the mid-season (July-August 2019) in the Prairie Coteau region near Brookings, South Dakota. Site abbreviations are as follows: DC = Deer Creek, AG = Altamont GPA, AR = Aurora, BP = Brookings Prairie, OL = Oak Lake Field Station, S7 = Seven-mile Fen, SV = Severson WMA, GG = Gary Gulch, CP = Coteau Prairie WMA, CL = Coteau Lakes GPA, RBH2 = Round Bullhead Lakes Site 2, SP = Sioux Prairie, AP = Altamont Prairie, RBH1 = Round Bullhead Site 1, J = Jacobsen Fen.

Fig. 6 Shannon diversity of biotically-pollinated plants at the family, genus, and species level during the late season (September-October 2019) in the Prairie Coteau region near Brookings, South Dakota. Prairies were only sampled once this season, thus are represented by a single bar in this graph. Site abbreviations are as follows: DC = Deer Creek, AG = Altamont GPA, AR = Aurora, BP = Brookings Prairie, OL = Oak Lake Field Station, S7 = Seven-mile Fen, SV = Severson WMA, GG = Gary Gulch, CP = Coteau Prairie WMA, CL = Coteau Lakes GPA, RBH2 = Round Bullhead Lakes Site 2, SP = Sioux Prairie, AP = Altamont Prairie, RBH1 = Round Bullhead Site 1, J = Jacobsen Fen.

#### Figure 1

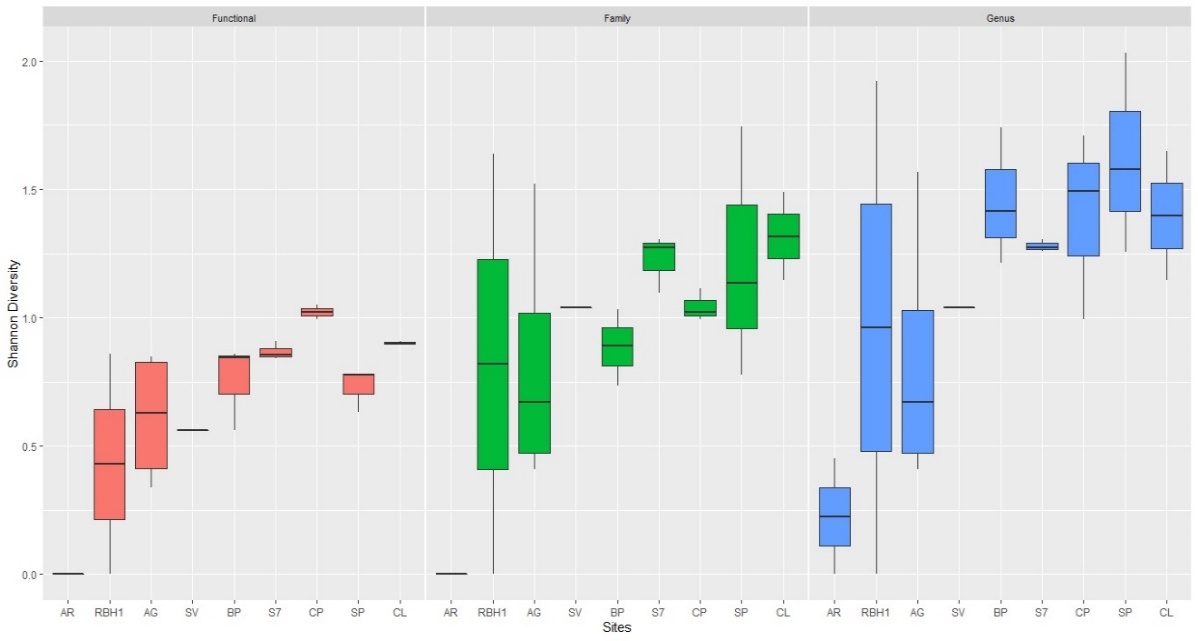

#### Figure 2

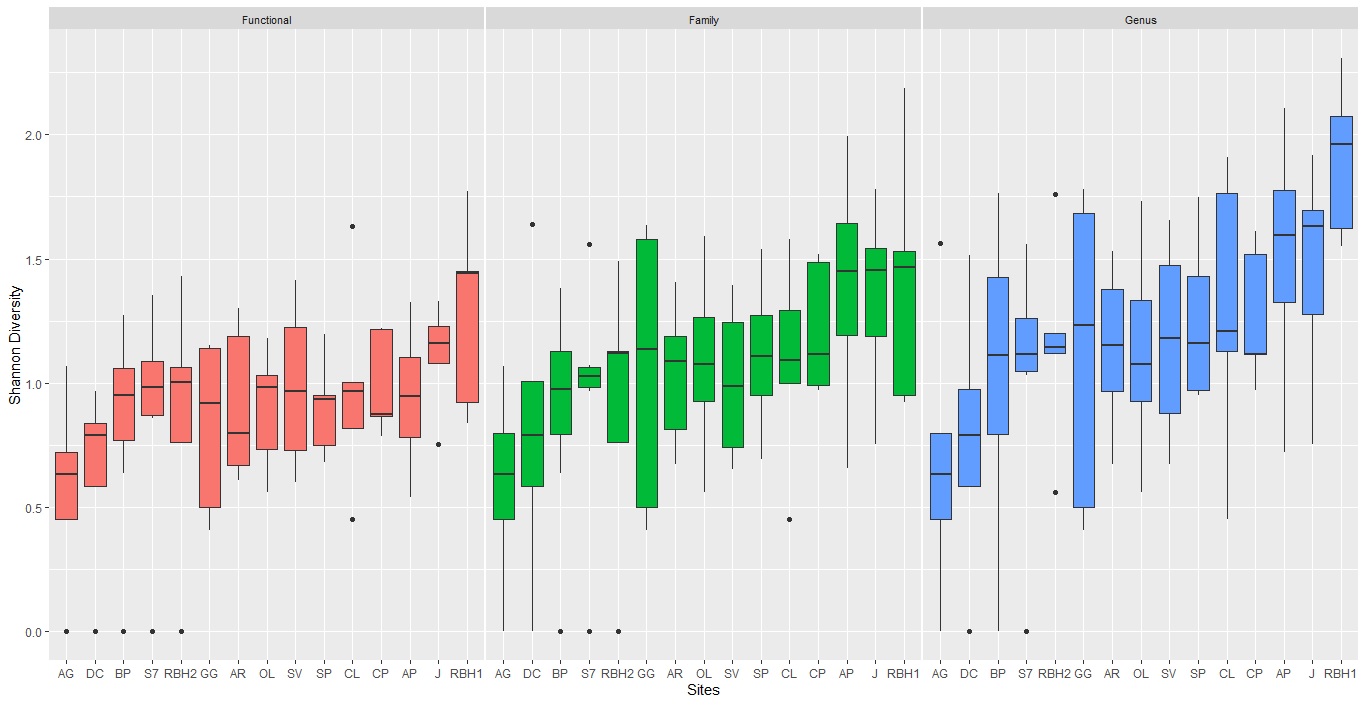

#### Figure 3

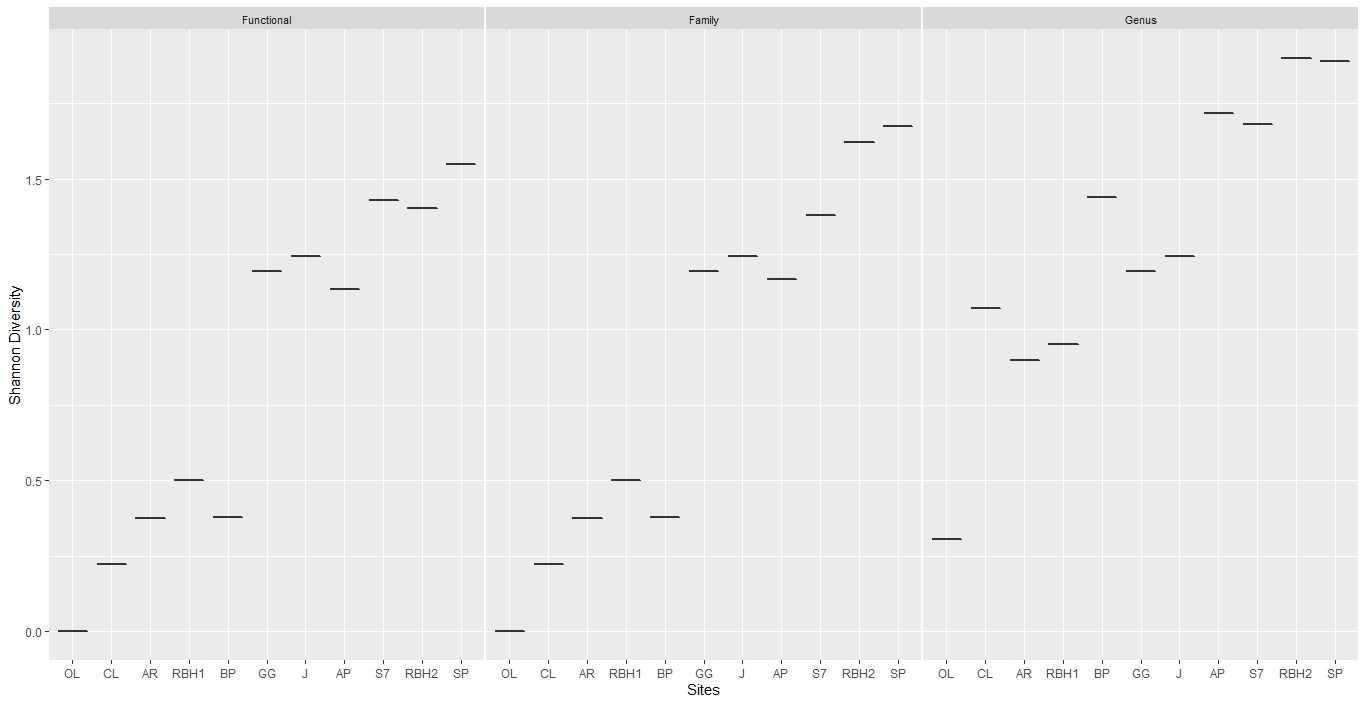

#### Figure 4

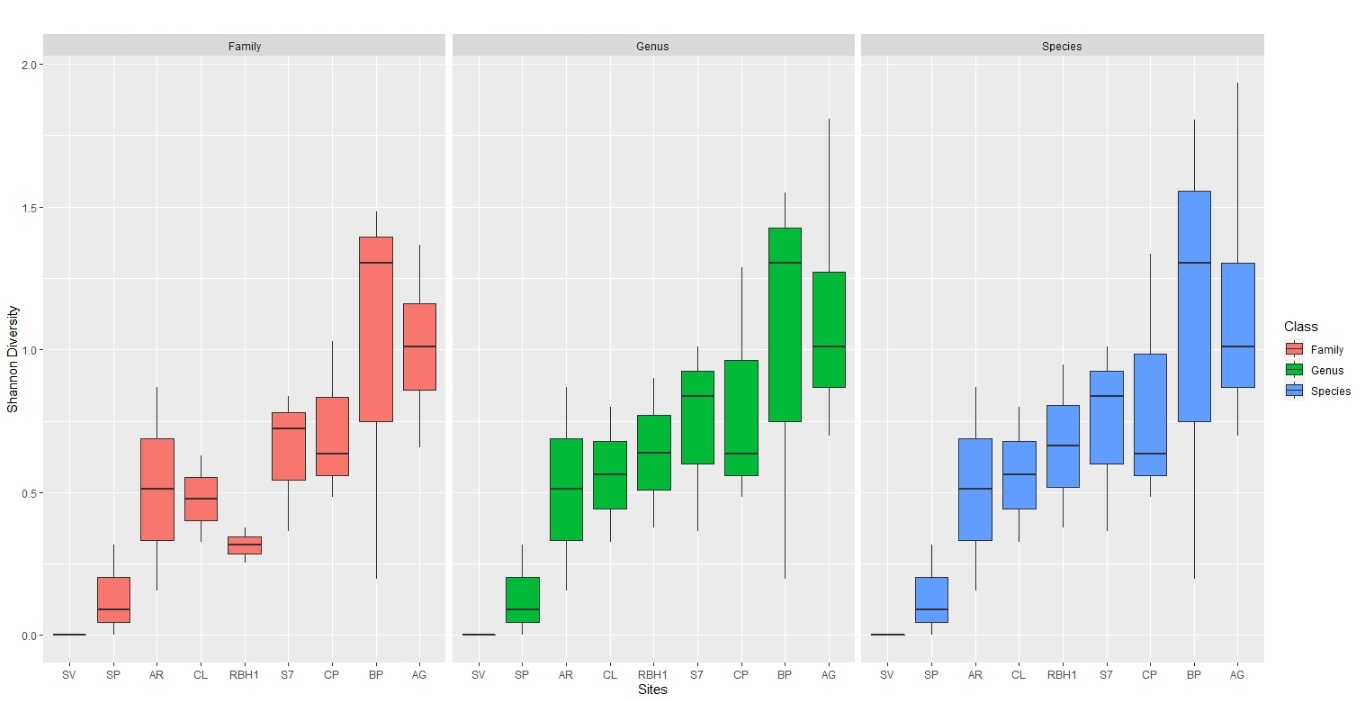

#### Figure 5

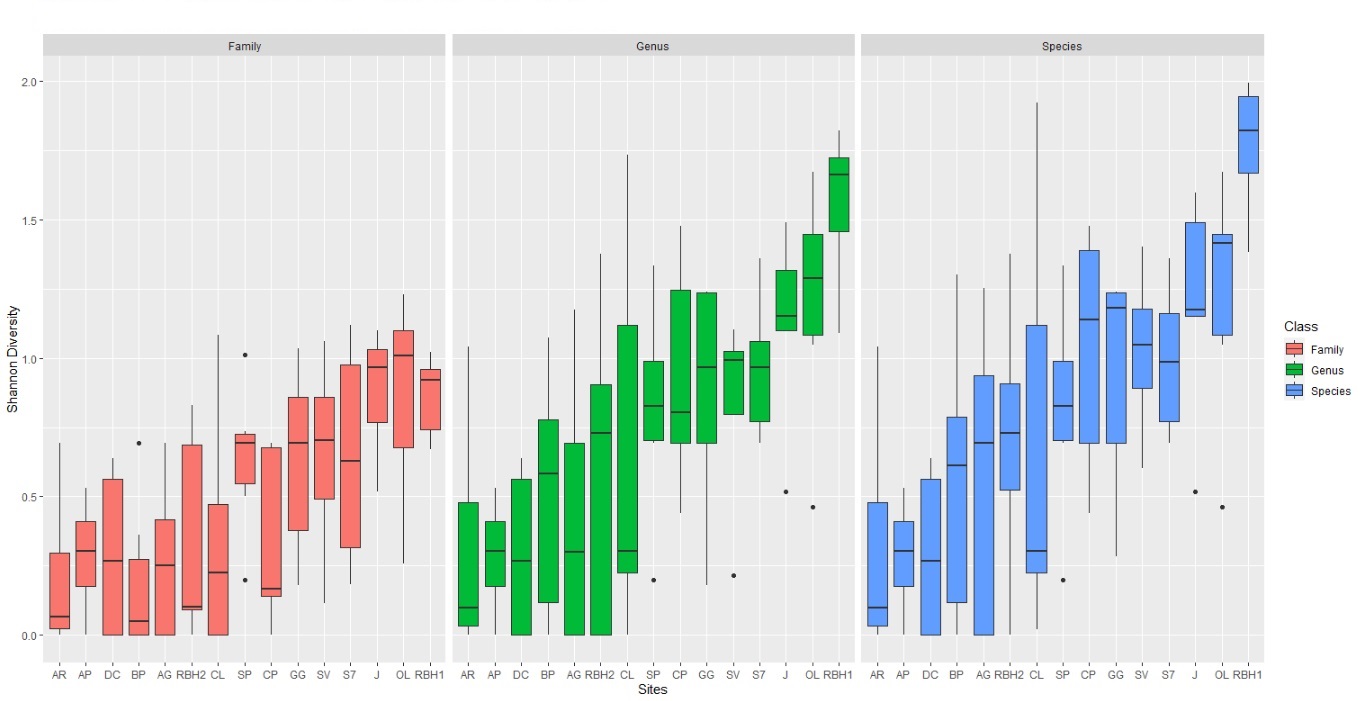

#### Figure 6

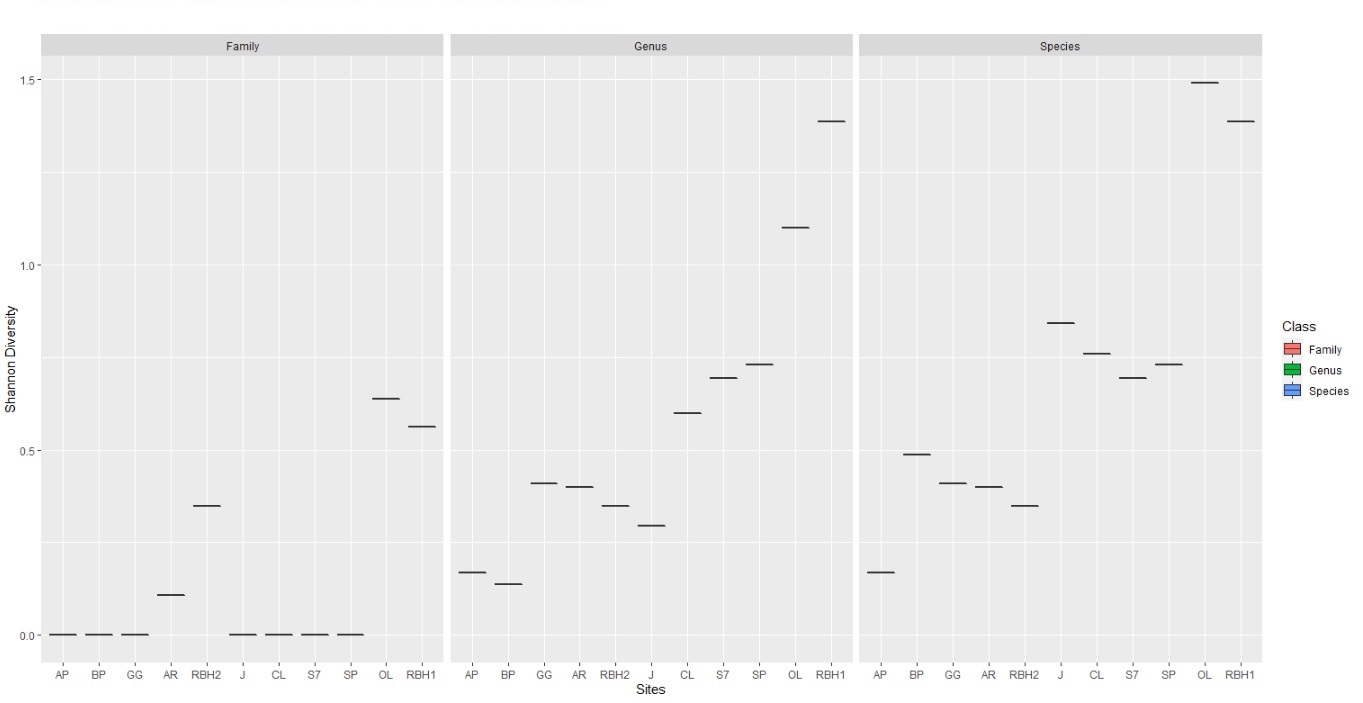

### Supplemental tables for: Quantifying habitat and landscape effects on composition and structure of plant-pollinator networks in the US Northern Great Plains

Table 1**.** Univariate generalized linear regression analysis of pollinator genus diversity on network-level specialization (H2’) for early (May-June), mid (July-August), and late (September-October) seasons in the Prairie Coteau region near Brookings, South Dakota in 2019. *P* < 0.05; * *P* < 0.05; ** *P* < 0.01; *** *P* < 0.001

| **Variable** | **Coefficient** | **Standard Error** | **Degrees of Freedom** | **p-value** |
| --- | --- | --- | --- | --- |
| Mid-season pollinator genus diversity  Intercept | 0.304  -0.01 | 0.072  0.14 | 12 | 0.001**  0.96 |
| Early season pollinator genus diversity  Intercept | -0.198  0.905 | 0.140  0.234 | 6 | 0.208  0.008** |
| Late season pollinator genus diversity  Intercept | 0.325  0.394 | 0.232  0.312 | 6 | 0.210  0.254 |

Table 2. Univariate generalized linear regression analysis of pollinator genus diversity on connectance for early (May-June), mid (July-August), and late (September-October) seasons in the Prairie Coteau region near Brookings, South Dakota in 2019. *P* < 0.05; * *P* < 0.05; ** *P* < 0.01; *** *P* < 0.001

| **Variable** | **Coefficient** | **Standard Error** | **Degrees of Freedom** | **p-value** |
| --- | --- | --- | --- | --- |
| Mid-season pollinator genus diversity  Intercept | -0.077  0.325 | 0.032  0.066 | 12 | 0.034*  0.0003*** |
| Early season pollinator genus diversity  Intercept | -0.258  0.775 | 0.106  0.178 | 6 | 0.052  0.0048** |
| Late season pollinator genus diversity  Intercept | -0.227  0.731 | 0.058  0.078 | 6 | 0.008**  8.58e-05*** |

Table 3. Univariate generalized linear regression analysis of pollinator genus diversity on nestedness for early (May-June), mid (July-August), and late (September-October) seasons in the Prairie Coteau region near Brookings, South Dakota in 2019. *P* < 0.05; * *P* < 0.05; ** *P* < 0.01; *** *P* < 0.001

| **Variable** | **Coefficient** | **Standard Error** | **Degrees of Freedom** | **p-value** |
| --- | --- | --- | --- | --- |
| Mid-season pollinator genus diversity  Intercept | -3.015  23.23 | 4.29  8.77 | 12 | 0.49  0.02* |
| Early season pollinator genus diversity  Intercept | 18.387  -10.101 | 3.488  5.827 | 6 | 0.002*  0.134 |
| Late season pollinator genus diversity  Intercept | 26.491  1.058 | 10.995  14.801 | 6 | 0.053  0.945 |

Table 4. Table of Spearman rank correlations for pollinator and plant diversities with Rho values and Bonferroni-corrected p values from early (May-June), mid (July-August), and late (September-October) seasons in the Prairie Coteau region near Brookings, South Dakota in 2019. *P* < 0.05; * *P* < 0.05; ** *P* < 0.01; *** *P* < 0.001

| **Season** | **Metrics tested** | **p-value** | **Rho value** |
| --- | --- | --- | --- |
| **Total** | Pollinator Functional Diversity & Pollinator Family Diversity | 7.92E-15 | 0.805 |
| **Total** | Pollinator Functional Diversity & Pollinator Genus Diversity | 6.98E-13 | 0.639 |
| **Total** | Pollinator Functional Diversity & Pollinator Species Diversity | 1.05E-05 | 0.458 |
| **Total** | Pollinator Family Diversity & Pollinator Genus Diversity | 2.27E-10 | 0.588 |
| **Total** | Pollinator Family Diversity & Pollinator Species Diversity | 4.57E-06 | 0.471 |
| **Total** | Pollinator Genus Diversity & Pollinator Species Diversity | 7.92E-15 | 0.749 |
| **Total** | Floral Family Diversity & Floral Genus Diversity | 7.92E-15 | 0.761 |
| **Total** | Floral Family Diversity & Floral Species Diversity | 7.92E-15 | 0.7 |
| **Total** | Floral Genus Diversity & Floral Species Diversity | 7.92E-15 | 0.968 |
| **Early** | Pollinator Functional Diversity & Pollinator Family Diversity | 7.92E-15 | 0.805 |
| **Early** | Pollinator Functional Diversity & Pollinator Genus Diversity | 6.98E-13 | 0.64 |
| **Early** | Pollinator Functional Diversity & Pollinator Species Diversity | 1.05E-05 | 0.458 |
| **Early** | Pollinator Family Diversity & Pollinator Genus Diversity | 2.27E-10 | 0.588 |
| **Early** | Pollinator Family Diversity & Pollinator Species Diversity | 4.57E-06 | 0.471 |
| **Early** | Pollinator Genus Diversity & Pollinator Species Diversity | 7.92E-15 | 0.75 |
| **Early** | Floral Family Diversity & Floral Genus Diversity | 7.92E-15 | 0.761 |
| **Early** | Floral Family Diversity & Floral Species Diversity | 7.92E-15 | 0.7 |
| **Early** | Floral Genus Diversity & Floral Species Diversity | 7.92E-15 | 0.968 |
| **Mid** | Pollinator Functional Diversity & Pollinator Family Diversity | 7.92E-15 | 0.805 |
| **Mid** | Pollinator Functional Diversity & Pollinator Genus Diversity | 6.98E-13 | 0.64 |
| **Mid** | Pollinator Functional Diversity & Pollinator Species Diversity | 1.05E-05 | 0.458 |
| **Mid** | Pollinator Family Diversity & Pollinator Genus Diversity | 2.27E-10 | 0.588 |
| **Mid** | Pollinator Family Diversity & Pollinator Species Diversity | 4.57E-06 | 0.471 |
| **Mid** | Pollinator Genus Diversity & Pollinator Species Diversity | 7.92E-15 | 0.75 |
| **Mid** | Floral Family Diversity & Floral Genus Diversity | 7.92E-15 | 0.761 |
| **Mid** | Floral Family Diversity & Floral Species Diversity | 7.92E-15 | 0.7 |
| **Mid** | Floral Genus Diversity & Floral Species Diversity | 7.92E-15 | 0.968 |
| **Late** | Pollinator Functional Diversity & Pollinator Family Diversity | 7.92E-15 | 0.805 |
| **Late** | Pollinator Functional Diversity & Pollinator Genus Diversity | 6.98E-13 | 0.64 |
| **Late** | Pollinator Functional Diversity & Pollinator Species Diversity | 1.05E-05 | 0.458 |
| **Late** | Pollinator Family Diversity & Pollinator Genus Diversity | 2.27E-10 | 0.588 |
| **Late** | Pollinator Family Diversity & Pollinator Species Diversity | 4.57E-06 | 0.471 |
| **Late** | Pollinator Genus Diversity & Pollinator Species Diversity | 7.92E-15 | 0.75 |
| **Late** | Floral Family Diversity & Floral Genus Diversity | 7.92E-15 | 0.761 |
| **Late** | Floral Family Diversity & Floral Species Diversity | 7.92E-15 | 0.7 |
| **Late** | Floral Genus Diversity & Floral Species Diversity | 7.92E-15 | 0.968 |
